# Fluorescence Activation Mechanism and Imaging of Drug Permeation with New Sensors for Smoking-Cessation Ligands

**DOI:** 10.1101/2021.10.04.463082

**Authors:** Aaron L. Nichols, Zack Blumenfeld, Chengcheng Fan, Laura Luebbert, Annet E. M. Blom, Bruce N. Cohen, Jonathan S. Marvin, Philip M. Borden, Charlene H. Kim, Anand K. Muthusamy, Amol V. Shivange, Hailey J. Knox, Hugo Rego Campello, Jonathan H. Wang, Dennis A. Dougherty, Loren L. Looger, Timothy Gallagher, Douglas C. Rees, Henry A. Lester

## Abstract

Nicotinic partial agonists provide an accepted aid for smoking cessation and thus contribute to decreasing tobacco-related disease. Improved drugs constitute a continued area of study. However, there remains no reductionist method to examine the cellular and subcellular pharmacokinetic properties of these compounds in living cells. Here, we developed new intensity-based drug sensing fluorescent reporters (“iDrugSnFRs”) for the nicotinic partial agonists dianicline, cytisine, and two cytisine derivatives – 10-fluorocytisine and 9-bromo-10-ethylcytisine. We report the first atomic-scale structures of liganded periplasmic binding protein-based biosensors, accelerating development of iDrugSnFRs and also explaining the activation mechanism. The nicotinic iDrugSnFRs detect their drug partners in solution, as well as at the plasma membrane (PM) and in the endoplasmic reticulum (ER) of cell lines and mouse hippocampal neurons. At the PM, the speed of solution changes limits the growth and decay rates of the fluorescence response in almost all cases. In contrast, we found that rates of membrane crossing differ among these nicotinic drugs by > 30 fold. The new nicotinic iDrugSnFRs provide insight into the real-time pharmacokinetic properties of nicotinic agonists and provide a methodology whereby iDrugSnFRs can inform both pharmaceutical neuroscience and addiction neuroscience.

## INTRODUCTION

Smoking cessation is an important goal to help decrease the burden, both individual and societal, of tobacco-related disease. The addictive tobacco alkaloid nicotine itself, via transdermal patches and other devices, remains available for people trying to quit smoking; but nicotine replacement therapy has distressingly low rates of success. Therefore, various research projects are continuing with the aim of developing more effective ligands for nicotinic acetylcholine receptors (nAChRs).

Prior work suggests that partial agonists with lower efficacy than nicotine could serve as effective smoking-cessation drugs (Rose, et al., 1994), and efforts continue in that direction (Rollema & Hurst, 2018). Another plant alkaloid, (-)-cytisine (also called cytisinicline and Tabex®), an α4β2 nAChR partial agonist, served as a basis for the synthesis of analogs which have not yet entered the clinic (Chellappan, Xiao, Tueckmantel, Kellar, & Kozikowski, 2006; Houllier, Gouault, Lasne, & Rouden, 2006; Imming, Klaperski, Stubbs, Seitz, & Gundisch, 2001; Kozikowski, et al., 2007; Marcaurelle, Johannes, Yohannes, Tillotson, & Mann, 2009; Philipova, et al., 2015; Rouden, et al., 2002). Varenicline (Chantix®) has four rings, two more than nicotine or cytisine, and is currently the only FDA-approved smoking-cessation drug, but the modest quit rate of ~18% at 12 months invites further investigation (Coe, et al., 2005; Mills, Wu, Spurden, Ebbert, & Wilson, 2009). Dianicline, another tetracyclic compound, was discontinued after unfavorable Phase III clinical trials (Cohen, et al., 2003; Fagerstrom & Balfour, 2006).

A nicotinic ligand for smoking cessation must satisfy at least three criteria (Rollema, et al., 2010; Tashkin, 2015). 1) It must enter the brain, where the most nicotine-sensitive nAChRs (α4β2) occur. It must also 2) activate α4β2 nAChRs with an EC_50_ sufficient to reduce cravings and withdrawal (1–2 μM). Finally, it must 3) block nicotine binding to reduce the reward phase of smoking (2 to 30 min). Varenicline meets these criteria, while cytisine (low brain penetration) and dianicline (EC_50_ =18 μM) each fail one of the criteria (Rollema, et al., 2010).

Membrane permeation is interesting for investigating and treating nicotine addiction in at least two ways. Firstly, note criterion #1 above. For uncharged molecules, the conventional metric for membrane permeability is logP, where P is the octanol-water partition coefficient. For weak bases including most orally available neural drugs, logP must be corrected to account for the fraction of uncharged (deprotonated) molecules at the pH of interest, usually pH7.4; the resulting metric, termed LogD_pH7.4_, is always less positive than logP. Enhancing the membrane permeability of cytisine analogs and probing nAChR subtype selectivity was addressed via direct functionalization of (-)-cytisine within the pyridone ring (Rego Campello, et al., 2018). Two of the resulting derivatives, 10-fluorocytisine and 9-bromo-10-ethylcytisine, have cytisine-like EC_50_ for the α4β2 nAChRs, but more positive calculated LogD_pH7.4_ values, suggesting greater membrane permeability at the nearly neutral pH of the blood, brain, and cytoplasm (Blom, Campello, Lester, Gallagher, & Dougherty, 2019). Estimates of LogD_pH7.4_ are inexact, extrapolated, or rely on algorithmic calculations whose results differ over two log units for individual molecules (Pienko, Grudzien, Taciak, & Mazurek, 2016). These estimates have unknown applicability to biological membranes at the LogD_pH7.4_ values < 0 that characterize varenicline, dianicline, and the cytisine analogs.

Secondly, nicotine dependence involves one or more “inside-out” mechanisms. Nicotine itself (logD_pH7.4_ 0.99) enters the endoplasmic reticulum (ER), binds to nascent nAChRs, becomes a pharmacological chaperone for the nAChRs, and eventually causes selective upregulation of these receptors on the plasma membrane (PM) (Henderson & Lester, 2015). For this reason, it is especially important to understand permeation into the ER.

These two neuroscience aspects of nicotinic ligands—pharmaceutical science and addiction science—call for direct measurements of drug movements in living cells. We previously explored the subcellular pharmacokinetics of nicotine and varenicline in immortalized cell lines and cultured neurons using the iDrugSnFRs iNicSnFR3a and iNicSnFR3b, to visualize that these nicotinic agonists enter the ER within seconds of drug application and exit equally rapidly from the ER upon extracellular washing (Shivange, et al., 2019). That nicotine diffuses across cellular membranes in seconds has been suspected for decades: nicotine crosses six plasma membranes to enter the brain within 20 s, providing a “buzz”. That varenicline becomes trapped in acidic vesicles suggests appreciable membrane permeation but may also underlie unwanted effects (Govind, et al., 2017; Le Houezec, 2003).

We sought to generate and to apply additional intensity-based drug-sensing fluorescent reporters (“iDrugSnFRs”) for candidate smoking cessation drugs: dianicline, cytisine, 10-fluorocytisine, and 9-bromo-10-ethylcytisine. We hypothesized that a family of newly developed iDrugSnFRs would enable quantifiable fluorescence signals that compare the differences in permeation among these compounds.

## RESULTS

### Generation of additional nicotinic iDrugSnFRs: structural tactic

To generate iDrugSnFRs for cytisine and dianicline, we followed two converging tactics. In the “structure-based” tactic, we obtained the first structural data for OpuBC-based SnFRs bound by nicotinic ligands (nicotine and varenicline) (Fig. 1, Supplementary Table 1). Crystals of iNicSnFR3adt in the presence of 10 mM nicotine diffracted to 2.95 Å resolution (PDB 7S7U). Overall, the liganded PBP domain of iNicSnFR3adt adopts a closed conformation (Fig. 1A). In the binding pocket between the top and bottom lobes of the PBP, we observed an “avocado”-shaped electron density in the nicotine binding site, enclosed by several aromatic residues (Fig. 1B). The combination of protonation/deprotonation and the rotatable bond of nicotine (Elmore & Dougherty, 2000) vitiate unambiguously localizing it within the binding pocket

**Figure 1.**
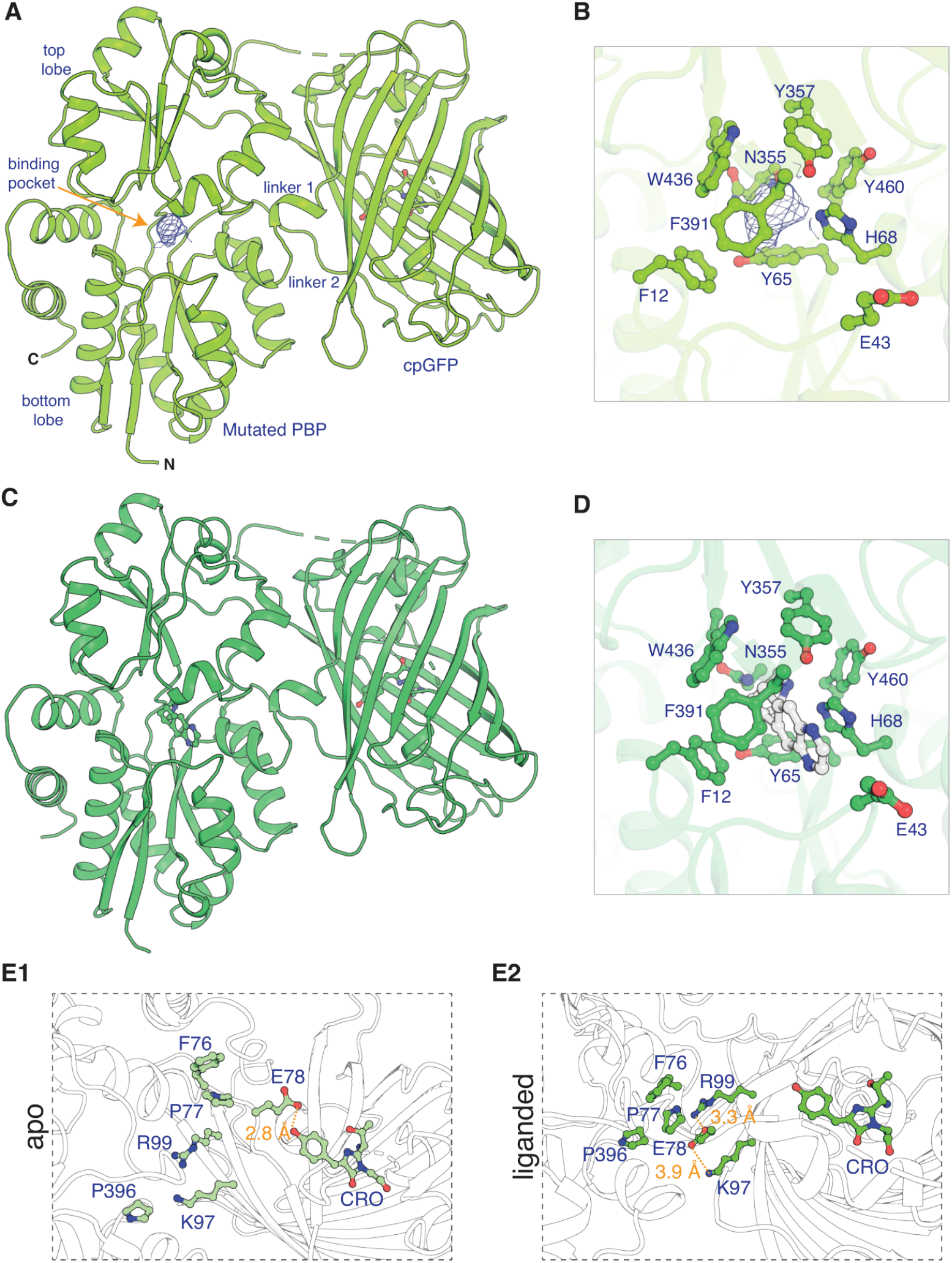
Apo and ligand-bound structures of iNicSnFR3adt (dt indicates that His_6_ and Myc tags have been removed to aid crystallization). To form an iDrugSnFR, a circularly permuted GFP molecule, flanked by two 4-residue linking sequences, is inserted into a PBP at a position (77-78, in our numbering system) that changes backbone Φ-Ψ angles between the apo and liganded PBP. **(A)** Overall conformation of iNicSnFR3adt crystallized with nicotine; an electron density appears at the nicotine binding site (PDB 7S7U). **(B)** iNicSnFR3adt binding site residues. **(C)** Overall conformation of iNicSnFR3adt with varenicline bound (PDB 7S7T). (**D**) iNicSnFR3adt binding site with varenicline present. **(E)** Aspects of the PBP-Linker1-cpGFP interface, emphasizing contacts that change upon ligand binding. The Phe76-Pro77-Glu78 cluster (in Linker 1) lies 11 to 16 Å from position 43, which defines the outer rim of the ligand site **(B)**; therefore, the cluster makes no direct contact with the ligand site. **(E1)** In the apo conformation, Glu78 acts as a candle snuffer that prevents fluorescence by the chromophore (PDB 7S7V). **(E2)** In the liganded conformation (PDB 7S7T), the Phe76-Pro77-Glu78 cluster moves Glu78 at least 14 Å away from the fluorophore. Pro77 is flanked by Phe76 and Pro396 (in the top lobe of the PBP moiety). The presumably deprotonated Glu78 forms salt bridges with Lys97 and Arg99, both facing outward on the β6 strand of the original GFP (within the original Phe165-Lys-Ile-Arg-His sequence).

We obtained an unambiguous ligand placement for iNicSnFR3adt in the presence of 10 mM varenicline in the same crystallization condition. Crystals of iNicSnFR3adt with varenicline bound were isomorphous to those of the nicotine-bound crystals and diffracted to 3.2 Å resolution (PDB 7S7T). While the protein structure (Fig. 1D) is identical to that of the nicotine bound structure (Fig. 1A), the rigidity and additional ring of varenicline allowed us to unambiguously localize it in the binding pocket. Varenicline is enclosed by the same aromatic residues as nicotine, forming cation-π interactions with Tyr65 and Tyr357, in addition to other interactions with the pocket residues (Fig. 1E).

The data confirm that similar ligand-induced conformational changes occur in the periplasmic binding protein (PBP) for nicotine, varenicline, ACh (Borden, et al., 2019), and choline (Fan, 2020) (Figure 1-figure supplement 1). These changes resemble those for other OpuBC PBPs (Schiefner, et al., 2004).

In the full iDrugSnFR, in the apo state, the Glu78 in linker 1 approaches within ~ 2.5 Å of the oxygen of the tyrosine fluorophore (Figure 1E1) (PDB 7S7V). Figure 1E2 provides structural details confirming the hypothesis (Barnett, Hughes, & Drobizhev, 2017; Nasu, Shen, Kramer, & Campbell, 2021) that in the liganded state, Glu78 has moved away, presumably allowing the fluorescent tyrosinate to form (Supplementary Video 1). We term this mechanism the “candle snuffer”.

### Generation of additional nicotinic iDrugSnFRs: mutational tactic

In the mutational tactic, we screened each drug shown in Figure 1-figure supplement 2 against a panel of biosensors that included iNicSnFR3a and iNicSnFR3b (Shivange, et al., 2019) and iAChSnFR (Borden, et al., 2019) as well as intermediate constructs from their development process. From this screen, we chose sensors with the lowest EC_50_ for each drug as our starting protein for iDrugSnFR evolution.

Because the candle snuffer mechanism explains several details of the agonist- and pH-sensitivity of both iNicSnFR3a and iSketSnFR (see Discussion), we presume that it represents a general mechanism for OpuBC-cpGFP SnFRs. We did not mutate residues that lie (in 3D space) between the binding site and linkers.

For dianicline and cytisine separately, we incrementally applied site-saturation mutagenesis (SSM) to first and second shell amino acid positions within the binding pocket. We evaluated each biosensor and drug partner in lysate from *E. coli* and carried forward the biosensor with the highest S-slope to the subsequent round. S-slope,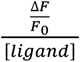at the beginning of the dose-response relation, emphasizes the response to ligand concentrations in the pharmacologically relevant range (Bera, et al., 2019). Table 1 and Fig. 2 summarize dose-response relations for the optimized sensors. The dianicline sensor, iDianiSnFR, has EC_50_ 6.7 ± 0.3 µM, ΔF_max_/F_0_ 7.4 ± 0.1, and S-slope 1.1. The cytisine sensor, iCytSnFR, has EC_50_ 9.4 ± 0.8 µM, ΔF_max_/F_0_ 5.0 ± 0.2, S-slope 0.5 (Table 1, Fig. 2A-B). After generating iCytSnFR, we performed additional SSM to progress from iCytSnFR to SnFRs for 10-fluorocytisine and 9-bromo-10-ethylcytisine. This optimization gave us iCyt_F_SnFR (EC_50_ 1.4 ± 0.04 µM, ΔF_max_/F0 7.9 ± 0.1, S-slope 5.6) and iCyt_BrEt_SnFR (EC_50_ 5.7 ± 0.1 µM, ΔF_max_/F0 4.0 ± 0.03, and S-slope 0.7) (Table 1, Fig. 2C-D).

**Table 1.**

| | | $\Delta F_{\max}/F_0$ | | $EC_{50}$ ($\mu M$) | | S-slope | | | | | | | | | |
| --- | --- | --- | --- | --- | --- | --- | --- | --- | --- | --- | --- | --- | --- | --- | --- |
| Informal Name | Drug of interest | L | P | L | P | L | P | 11 | 43 | 44 | 68 | 32<br>4 | 36<br>0 | 39<br>1 | 39<br>5 |
| iNicSnFR3b | nicotine | ND | 10 | ND | 19 | ND | 0.5 | E | E | N | H | S | T | F | G |
| iDianiSnFR | dianicline | $7.4 \pm 0.1$ | $4.7 \pm 0.2$ | $6.7 \pm 0.3$ | $15 \pm 1$ | 1.1 | 0.3 | D | R | - | S | N | G | - | N |
| iAChSnFR | ACh | ND | 12 | ND | 1.3 | ND | 9.2 | I | V | N | H | A | T | F | G |
| iCytSnFR | cytisine | $5.0 \pm 0.2$ | $7.3 \pm 0.4$ | $9.4 \pm 0.8$ | $11 \pm 1$ | 0.5 | 0.7 | - | Y | - | - | - | - | W | - |
| iCyt_F_SnFR | 10-fluorocytisine | $7.9 \pm 0.1$ | $2.3 \pm 0.1$ | $1.4 \pm 0.04$ | $1.6 \pm 0.3$ | 5.6 | 1.4 | - | N | G | - | - | - | W | - |
| iCyt_BrEt_SnFR | 9-bromo-10-ethylcytisine | $4.0 \pm 0.03$ | $3.6 \pm 0.04$ | $5.7 \pm 0.1$ | $4.2 \pm 0.2$ | 0.7 | 0.9 | - | Q | G | - | - | - | W | - |

**Nicotinic agonist iDrugSnFR naming, dose-response relations, and residues mutated.** Measurements in *E. coli* lysates (L) or with purified protein (P). ND, Not determined. Data for iAChSnFR from (Borden, et al., 2019); Data for iNicSnFR3b from (Shivange, et al., 2019).

**Figure 2.**
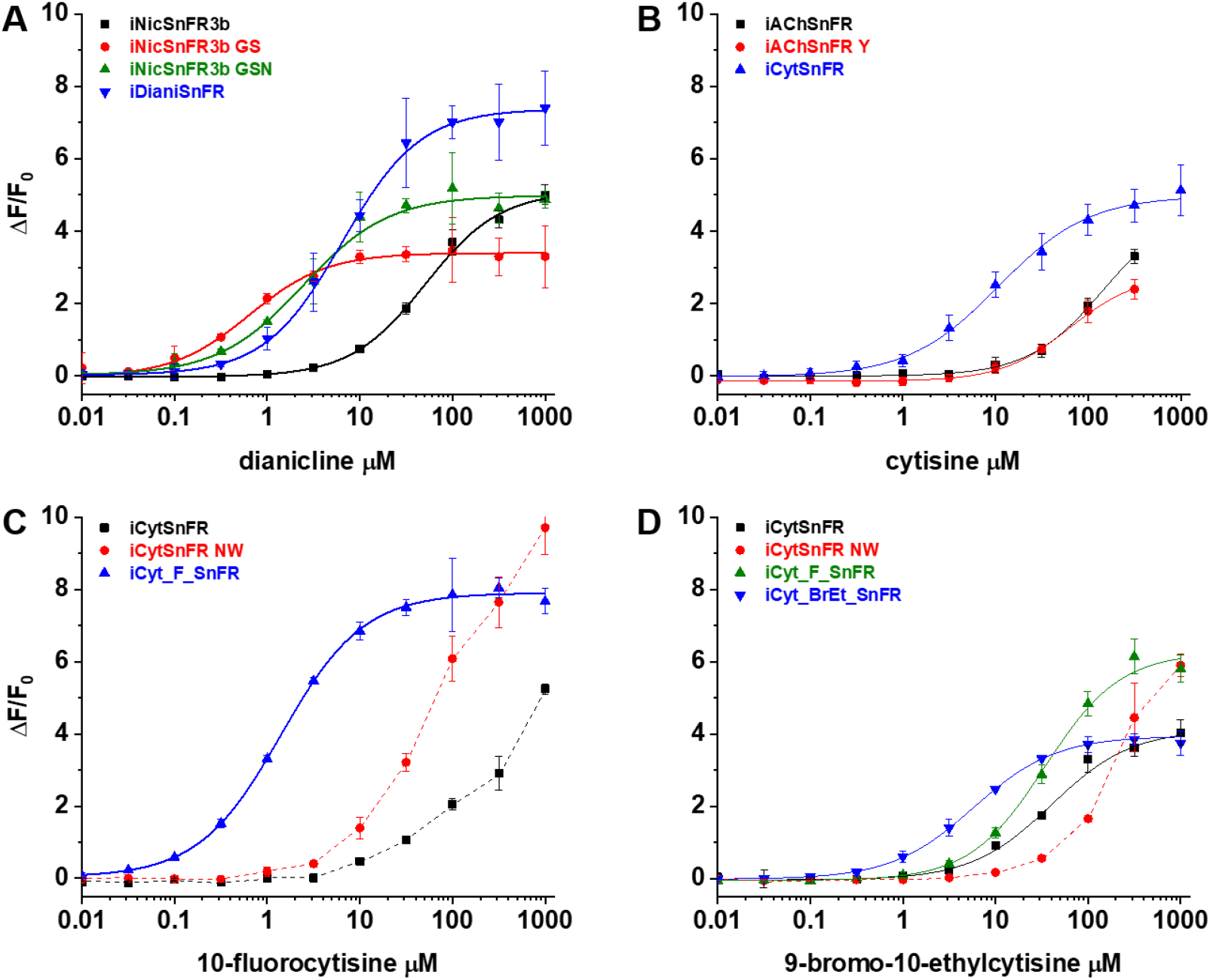
Nicotinic agonist iDrugSnFR development. Dose-response relations on intermediate constructs using *E. coli* lysate were performed with respective drug partners to identify SSM winners. **(A-D)** The progenitor biosensor is listed in black. Dashed lines indicate data that did not reach saturation at the concentrations tested; therefore, EC_50_ and ΔF_max_/F0 could not be determined. Development of **(A)** iDianiSnFR, **(B)** iCytSnFR, **(C)** iCyt_F_SnFR, and **(D)** iCyt_BrEt_SnFR.

### Specificity and thermodynamics of nicotinic iDrugSnFRs

We characterized the specificity of purified iDrugSnFRs for their drug partners versus a panel of related nicotinic agonists (Table 2, Fig. 3). The newly developed iDrugSnFRs showed some sensitivity to related nicotinic agonists. iDianiSnFR had the greatest fidelity for its drug partner but also showed an increased EC_50_ (15 µM) as a purified protein versus its EC_50_ in lysate (6.7 µM), possibly indicating decreased stability in a purified form. iCytSnFR, iCyt_F_SnFR, and iCyt_BrEt_SnFR showed a greater level of promiscuity for the compounds comprising the nicotinic agonist panel. Of note, iCytSnFR, iCyt_F_SnFR, and iCyt_BrEt_SnFR have an exceptionally low (60-90 nM) EC_50_ for varenicline. The newly developed iDrugSnFRs showed negligible binding to choline or the neurotransmitter acetylcholine, leading one to expect minimal endogenous interference during future *in vivo* experiments.

**Table 2.**

| Drug Name | iDianiSnFR |  |  | iCytSnFR |  |  | iCyt_F_SnFR |  |  | iCyt_BrEt_SnFR |  |  |
| --- | --- | --- | --- | --- | --- | --- | --- | --- | --- | --- | --- | --- |
| | $\Delta F_{\max}/F_0$ | EC<sub>50</sub> (μM) | S-slope | $\Delta F_{\max}/F_0$ | EC<sub>50</sub> (μM) | S-slope | $\Delta F_{\max}/F_0$ | EC<sub>50</sub> (μM) | S-slope | $\Delta F_{\max}/F_0$ | EC<sub>50</sub> (μM) | S-slope |
| choline | 2.0 ± 0.1 | 84 ± 20 | <0.1 | 5.8 ± 0.2 | 240 ± 30 | <0.1 | 2.6 ± 0.1 | 18 ± 1 | 0.1 | 2.6 ± 0.1 | 12 ± 1 | 0.2 |
| acetylcholine | 7.4 ± 1.0 | 660 ± 80 | <0.1 | 2.9 ± 0.1 | 35 ± 3 | <0.1 | 4.4 ± 0.3 | 222 ± 50 | <0.1 | 2.5 ± 0.2 | 73 ± 6 | <0.1 |
| cytisine | - | - | <0.1* | 7.3 ± 0.4 | 11 ± 1 | 0.7 | 4.4 ± 0.1 | 2.6 ± 0.3 | 1.7 | 4.7 ± 0.1 | 3.5 ± 0.2 | 1.3 |
| dianicline | 4.7 ± 0.2 | 15 ± 1 | 0.3 | 6.5 ± 0.4 | 34 ± 4 | 0.2 | 2.3 ± 0.3 | 43 ± 6 | <0.1 | 4-6 | >100 | <0.1** |
| nicotine | 2.2 ± 0.1 | 440 ± 100 | <0.1 | 6.4 ± 0.2 | 14 ± 2 | 0.5 | 4.7 ± 0.1 | 3.8 ± 0.2 | 1.2 | 4.8 ± 0.1 | 5.5 ± 0.2 | 0.9 |
| varenicline | 2.4 ± 2.0 | 1200 ± 500 | <0.1 | 6.5 ± 0.1 | 0.06 ± 0.01 | 110 | 7.1 ± 0.2 | 0.09 ± 0.02 | 79 | 5.3 ± 0.1 | 0.06 ± 0.01 | 88 |
| 10-fluorocytisine | ND | ND | ND | ND | ND | ND | 2.3 ± 0.1 | 1.6 ± 0.3 | 1.4 | 3.0 ± 0.1 | 4.7 ± 0.3 | 0.6 |
| 9-bromo-10-ethylcytisine | ND | ND | ND | ND | ND | ND | 3.1 ± 0.1 | 31 ± 2 | 0.1 | 3.6 ± <0.1 | 4.2 ± 0.2 | 0.9 |

**Figure 3.**
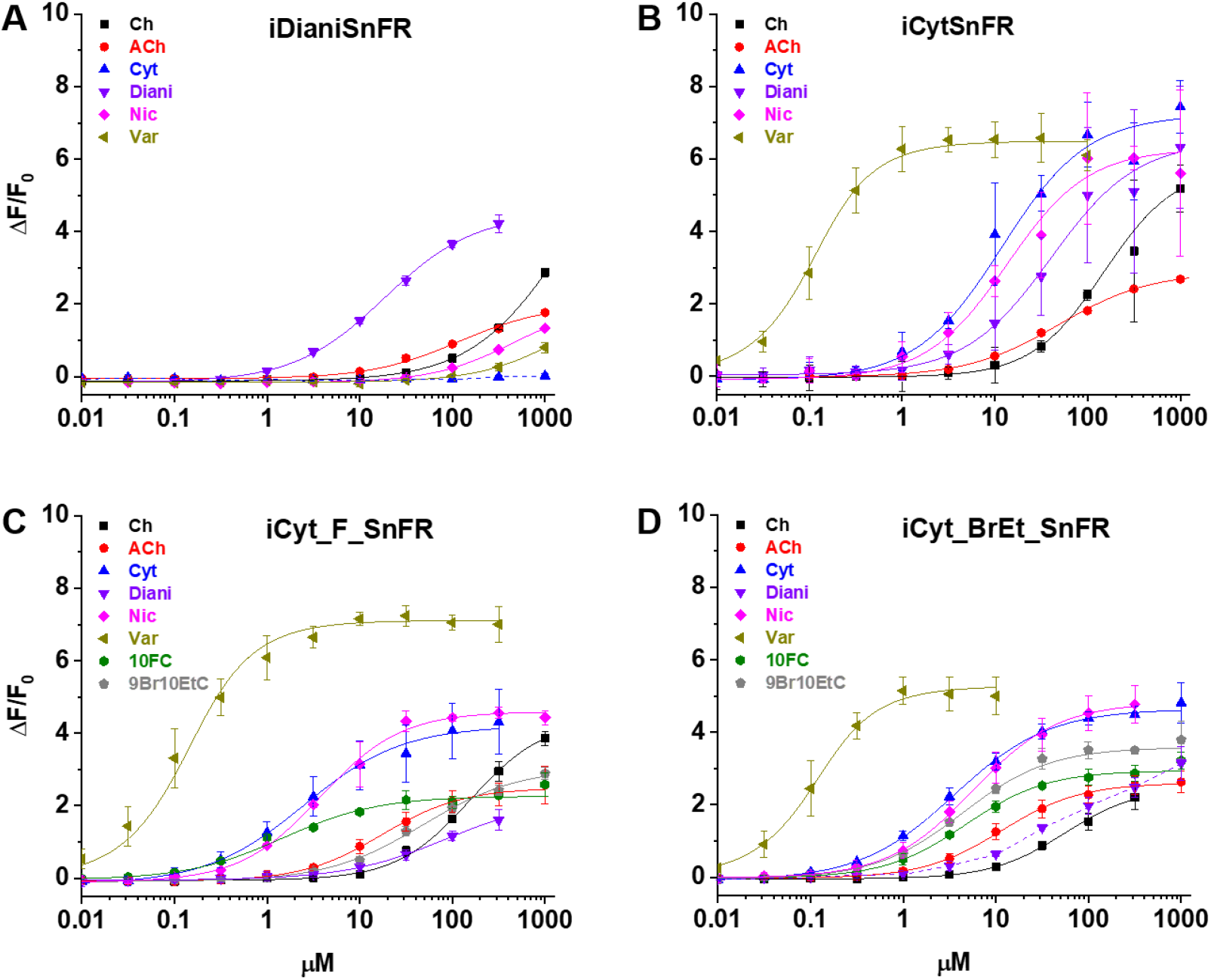
Dose-response relations of iDrugSnFR protein versus a nicotinic agonist panel. Abbreviations: Ch (choline), ACh (acetylcholine), Cyt (cytisine), Diani (dianicline), Nic (nicotine), Var (varenicline), 10FC (10-fluorocytisine), and 9Br10EtC (9-bromo-10-ethyl-cytisine). **(A-D)** Relevant EC_50_ values for each iDrugSnFR are listed in Table 2. Dashed lines indicate dose-response relations that did not approach saturation for the concentration ranges tested; therefore, EC_50_ and ΔF_max_/F0 could not be determined. **(A)** iDianiSnFR shows preference for dianicline, with some promiscuity for other nicotinic agonists. **(B)** iCytSnFR, **(C)** iCyt_F_SnFR, and **(D)** iCyt_BrEt_SnFR bind their drug partner, but also respond to other nicotinic agonists.

We also performed dose-response experiments with iDianiSnFR, iCytSnFR, iCyt_F_SnFR, and iCyt_BrEt_SnFR against a panel of nine endogenous molecules, including neurotransmitters (Supplementary Fig. 1). iDianiSnFR showed no response to any of the nine selected compounds above background. iCytSnFR, iCyt_F_SnFR, and iCyt_BrEt_SnFR showed no response above background for seven of the compounds. However, they exhibited a ΔF/F_0_ of 0.25-0.8 to dopamine at 316 µM/1 mM and a ΔF/F_0_ of 0.8–1.5 to serotonin (5-HT) at 316 µM/1 mM. In terms of S-slope, the relevant metric for most cellular or *in vivo* experiments, the SnFRs are at least 250-fold more sensitive to their eponymous partners than to other molecules we have tested.

To examine the thermodynamics of the iDrugSnFR:drug interaction, we conducted isothermal titration calorimetry (ITC) binding experiments (Fig. 4). The experimentally determined K_D_ of each iDrugSnFR:drug pair using ITC was within a factor of 1.5 from the experimentally determined EC_50_ for fluorescence in *E. coli* lysate or purified protein (Table 3). We infer that the EC_50_ for fluorescence is dominated by the overall binding of the ligand for all the iDrugSnFRs.

**Table 3.**

| **Biosensor** | **K<sub>D</sub> (μM)** | **n** | **ΔH**<br>**(kcal/mol)** | **-TΔS**<br>**(kcal/mol)** | **ΔG**<br>**(kcal/mol)** |
| --- | --- | --- | --- | --- | --- |
| **iCytSnFR** | **13.7 ± 1.1** | **0.84 ± 0.05** | **-2.1 ± 0.1** | **-4.6 ± 0.2** | **-6.6 ± 0.1** |
| **iCyt_F_SnFR** | **1.8 ± 0.5** | **0.83 ± 0.02** | **-5.5 ± 0.1** | **-2.4 ± 0.2** | **-7.9 ± 0.1** |
| **iCyt_BrEt_SnFR** | **5.4 ± 0.8** | **0.69 ± 0.09** | **-1.12 ± 0.03** | **6.1 ± 0.1** | **-7.2 ± 0.1** |
| **iDianiSnFR** | **7.6 ± 1.4** | **0.92 ± 0.02** | **3.2 ± 0.5** | **10.1 ± 0.4** | **-7.0 ± 0.2** |

**Affinity, occupancy number, and thermodynamic data calculated from isothermal titration calorimetry.** Data are the mean ± SEM, 3 runs.

**Figure 4.**
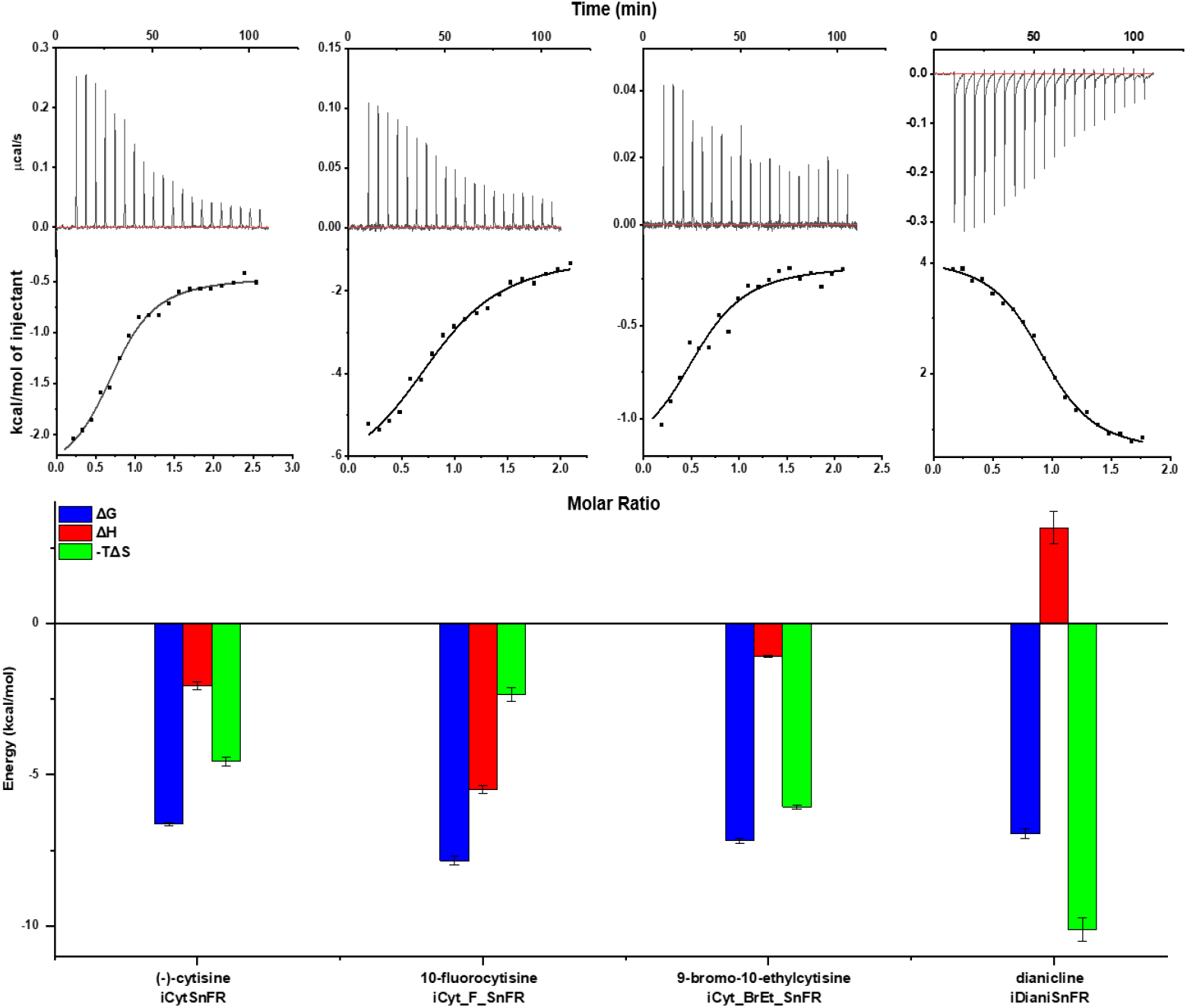
Isothermal titration calorimetry traces, fits, and thermodynamic data. **Top row**: Exemplar heat traces of iCytSnFR, iCyt_F_SnFR, iCyt_BrEt_SnFR, and iDianiSnFR paired with their drug partners obtained by isothermal calorimetry. The heats for iCytSnFR, iCyt_F_SnFR, and iCyt_BrEt_SnFR were exothermic, while that for iDianiSnFR was endothermic. **Middle row**: The resulting fits for each iDrugSnFR:drug pair from the integrated heats comprising each series of injections. **Bottom row:** Energy calculations. All iDrugSnFRs show exergonic reactions, but the relative enthalpic and entropic contributions vary among iDrugSnFRs. Data are from 3 separate runs, Mean ± SEM.

### Kinetics of nicotinic agonist iDrugSnFRs: stopped-flow

In a stopped-flow apparatus, we measured the fluorescence changes of iDrugSnFRs with millisecond resolution during both multiple 1 s trials and an independent 100 s trial. The stopped-flow data revealed that iDrugSnFRs do not have pseudo-first-order kinetic behaviors typical of two state binding interactions. Time courses of iDianiSnFR (both over 1 s and 100 s) were best fitted by double exponential equations. Most of the fluorescence change occurs within the first 0.1 s of mixing (Fig. 5A), with only minor additional increase by 100 s.

**Figure 5.**
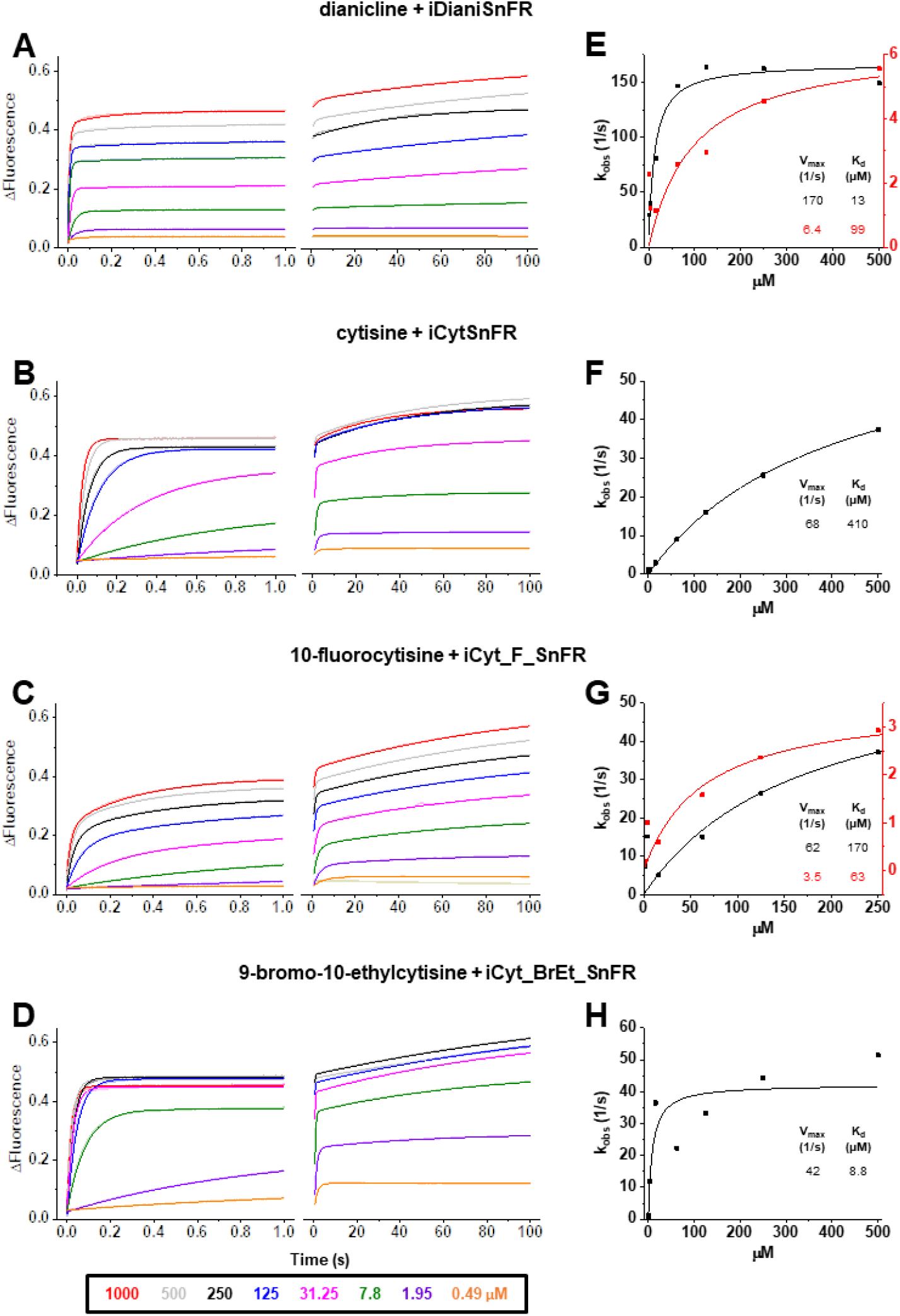
Stopped-flow fluorescence kinetic data for **(A)** iDianiSnFR, **(B)** iCytSnFR, **(C)** iCyt_F_SnFR, and **(D)** iCyt_BrEt_SnFR over 1 s and 100 s. Fluorescence was activated by mixing with the agonists as noted. Stopped-flow data shows a departure from first-order kinetics for this set of iDrugSnFRs. iDianiSnFR and iCyt_F_SnFR are fit to a double exponential; iCytSnFR and iCyt_BrEt_SnFR are fit to a single exponential. **(E-H)** Plots of the observed apparent rate constant against [agonist] for the 1 s data obtained in **(A-D)**. In H, we have confidence that the *k*_*obs*_ shows a maximal value of 40-50 s^-1^; the *K*_*d*_ probably lies within 2-fold of the fitted value.

Changes in fluorescence from iCytSnFR during the first 1 s of mixing fit well to a single exponential (Fig. 5B), and have close to pseudo-first order kinetics (i.e. the observed rate of fluorescence change is nearly linear with drug concentration). As with iDianiSnFR, most of the fluorescence change occurs within the first second, with additional fluorescent increase continuing over the next minute (Fig. 5B, right panel).

Like iDianiSnFR, iCyt_F_SnFR fluorescence changes are best fit by a double exponential (Fig. 5C), but the time course of fluorescence change is significantly slower. Fluorescence gradually increases throughout the recording period and beyond. This information was considered in later *in vitro* and *ex vivo* experiments.

iCyt_BrEt_SnFR fits well to a single exponential (Fig. 5D) for the first 1 s of data collection, but like the other sensors, continues to increase its fluorescence over longer periods.

We plotted the k_obs_ (s^-1^) obtained in the 1 s stopped-flow experiments versus concentration (Fig. 5E-5H). The aberrations from an ideal first-order kinetics vitiate generation of definitive k_off_ and k_on_ values (Supplementary Table 2), but we can approximate a K_max_ and K_D_ from our fitting procedures. Our stopped-flow experiments reinforced previous observations (Unger, et al., 2020) that the kinetics of iDrugSnFR binding involve complexities beyond a simple first-order kinetic model governing two binding partners.

### Kinetics of nicotinic agonist iDrugSnFRs: millisecond microperfusion

We also studied iCytSnFR_PM expressed in HEK293T cells during fluorescence responses to ACh, cytisine, or varenicline in a microperfusion apparatus that exchanged solutions near the cell on a millisecond time scale (Methods). This system directly measures the decay of the response when ligand is suddenly removed. The rank order of the iCytSnFR steady-state sensitivities is varenicline > cytisine > ACh. The time constant for decay decreased with increasing steady-state EC_50_ of the ligands, as though more tightly binding ligands dissociate more slowly (Fig. 6A).

**Figure 6.**
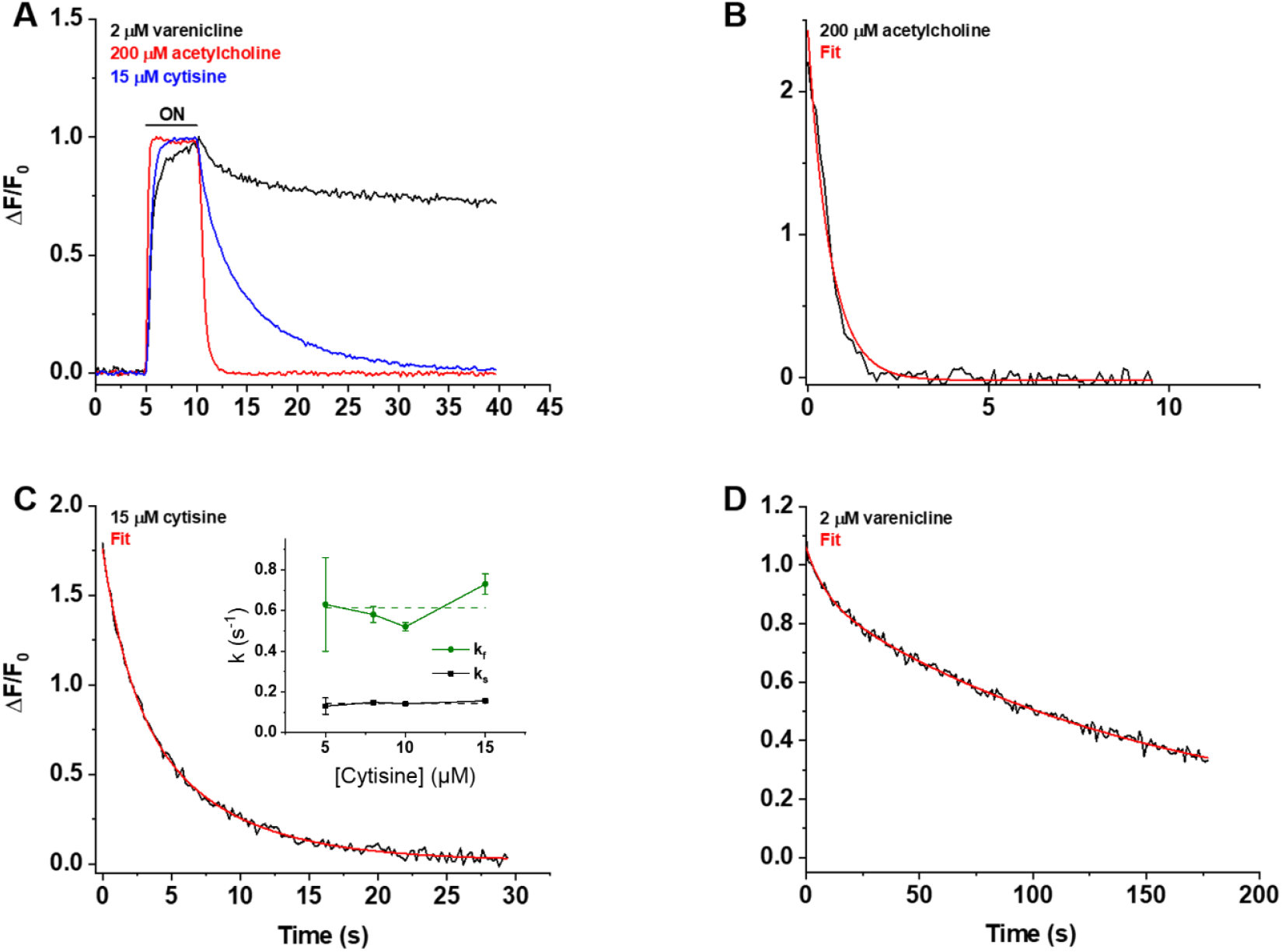
Decay of the iCytSnFR_PM responses after removal of ACh, cytisine, or varenicline. **(A)** The red, blue, and black traces are mean *ΔF/F*_*0*_ values for the ACh (200 µM), cytisine (15 µM), and varenicline (2 µM) responses as a function of time (n = 4-10 areas per ligand). The *ΔF/F*_*0*_ was normalized to the peak response for each ligand. Sampling rate was 5 frames/s. Ligand was applied for 5 s, denoted by the black horizontal bar above the traces. **(B-D)** Examples of the decay phase of the response to ACh (200 µM), cytisine (15 µM), and varenicline (2 µM) in individual areas (black traces in each panel). Red lines are fits to the sum of one or two negative exponential terms and a constant (red lines in each panel) using non-linear least-squares regression. **(B)** The decay of the ACh (200 µM) response (n = 1 area, 3 cells) was monophasic with a single time constant (*τ*_*0ff*_) of 0.61 ± 0.02 s (± SE, n = 86 frames, sampling rate of 9.8 frames/s). The red line is a fit to the sum of a negative exponential component (*R*^*2*^ of 0.98). **(C)** The decay of the cytisine (15 µM) response (n = 1 area, 4 cells) was biphasic with time constants (*τf*_*off*_, *τs*_*off*_) of 1.9 ± 0.2 and 6.6 ± 0.5 s (n = 149 frames, sampling rate of 5 frames/s). The red line is a fit to the sum of two negative exponential components and a constant (*R*^*2*^ of 0.996). It was significantly better than that of the sum of a single negative exponential term and a constant (F-test, *p* < 0.05). The relative amplitude of the slower decay component (*A*_*s*_*/(A*_*s*_*+A*_*f*_); where *A*_*s*_ is amplitude of the slower component of decay in units of *ΔF/F*_*0*_ and *A*_*f*_ is amplitude of the faster component) was 61%. Inset, neither rate constant changed significantly over the [cytisine] range from 5 to 15 μM. Dashed lines give the average over this range. **(D)** The decay of the varenicline (2 µM) response (n = 1 area, 3 cells) was also biphasic with a *τf*_*off*_ and *τs*_*off*_ of 9 ± 1 s and 150 ± 10 s (n = 178 frames, sampling rate of 1 frame/s), respectively. The *A*_*s*_*/(A*_*s*_*+A*_*f*_) was 83%. The red line is a fit to the sum of two negative exponential terms and a constant (*R*^*2*^ of 0.994) and it was significantly better than that to the sum of a single negative exponential term and a constant (F-test, *p* < 0.05).

We measured the decay waveforms after drug pulses at concentrations ≥ the EC_50_ of the steady-state response to maximize the *ΔF/F*_*0*_ signal/noise ratio (Fig. 6A-D). Because the decay phases are measured in zero [ligand], one expects that the decay rate constant(s) (*k*_*off*_) for an iDrugSnFR do not depend on the pulsed ligand concentration. Decay of the ACh response followed a single exponential time course (Fig. 6B). The values of the *k*_*off*_ for 30, 100, and 200 µM ACh did not differ significantly (ANOVA, *p* = 0.62, degrees of freedom (*df*) = 2 (model), 20 (error)). We pooled them to obtain a mean *k*_*off*_ of 1.9 ± 0.1 s^-1^ (mean ± SEM, n = 23 areas (50 cells)). The corresponding time constant *τ*_*0ff*_ was 530 ± 30 ms. Hence, the temporal resolution of the CytSnFR_PM sensor for changes in the ACh concentration was in the sub-second range.

The decay of the cytisine and varenicline response was biphasic (Fig. 6C-D): two exponential decay terms with an additional constant component fitted the cytisine decay significantly better than a single exponential term (F-test, *p* < 0.05). As expected, neither the faster decay rate constants (*kf*_*off*_) (ANOVA, *p* = 0.30, *df* = 3,32) nor the slower decay rate constants (*ks*_*off*_) (ANOVA, *p* = 0.54, *df* = 3,31) differed among the tested cytisine concentrations (5-15 µM). The *kf*_*off*_ and *ks*_*off*_ for 5-15 µM cytisine were 0.61 ± 0.04 s^-1^ (n = 36 areas, 105 cells) and 0.146 ± 0.006 s^-1^ (n = 35 areas, n = 103 cells), respectively. The corresponding decay time constants (*τf*_*0ff*_, *τs*_*0ff*_) were 1.8 ± 0.1 s and 6.9 ± 0.2 s. Therefore, the temporal resolution of CytSnFR_PM sensor for cytisine was < 10 s, adequate for the temporal resolution of the live-cell experiments presented below.

Interestingly, the decay waveform of the varenicline response was much slower than that for cytisine or ACh (Fig. 6A, 6D). We pulsed 2 µM varenicline, >> the EC_50_ of the steady-state response of the isolated protein (60 ± 10 nM) (Fig. 6D). The values of the *kf*_*off*_ and *ks*_*off*_ were 0.9 ± 0.2 s^-1^ and 0.0065 ± 0.0002 s^-1^, respectively (n= 4 areas (9 cells)). The slower component dominated the decay phase, with a fractional amplitude of 85 ± 1%. Thus, the temporal resolution of the iCytSnFR_PM sensor for varenicline was in the minute range. In the live-cell experiments described below, it would not be possible to resolve the differences between varenicline at the PM and in the ER. The relatively high affinity of iCytisineSnFR for varenicline, which presumably arises in part from the increased lifetime of the varenicline-iDrugSnFR complex, has drawbacks. The temporal resolution of iNicSnFR3a and iNicSnFR3b, which bind varenicline ~ 100-fold less tightly, is appropriate for subcellular experiments (Shivange, et al., 2019). The previous experiments showing ER entry of varenicline used iNicSnFR3a and iNicSnFR3b (Shivange, et al., 2019). For additional microperfusion data and analyses see Supplementary Figures 2-4.

### Characterization of nicotinic iDrugSnFRS in HeLa cells and primary mouse hippocampal culture

We examined the subcellular pharmacokinetics of the nicotinic agonists in mammalian cell lines and primary mouse hippocampal neurons. The nicotinic iDrugSnFRs were targeted to the plasma membrane (PM) (iDrugSnFR_PM) or the endoplasmic reticulum (ER) (iDrugSnFR_ER) as previously described (Bera, et al., 2019; Shivange, et al., 2019). We then performed a dose-response experiment using wide-field fluorescence imaging with each iDrugSnFR and its drug partner, sampling a range of concentrations covering a log scale surrounding the EC_50_ as determined for the purified protein (Fig. 7, Fig. 8, Supplementary Videos 2-5).

**Figure 7.**
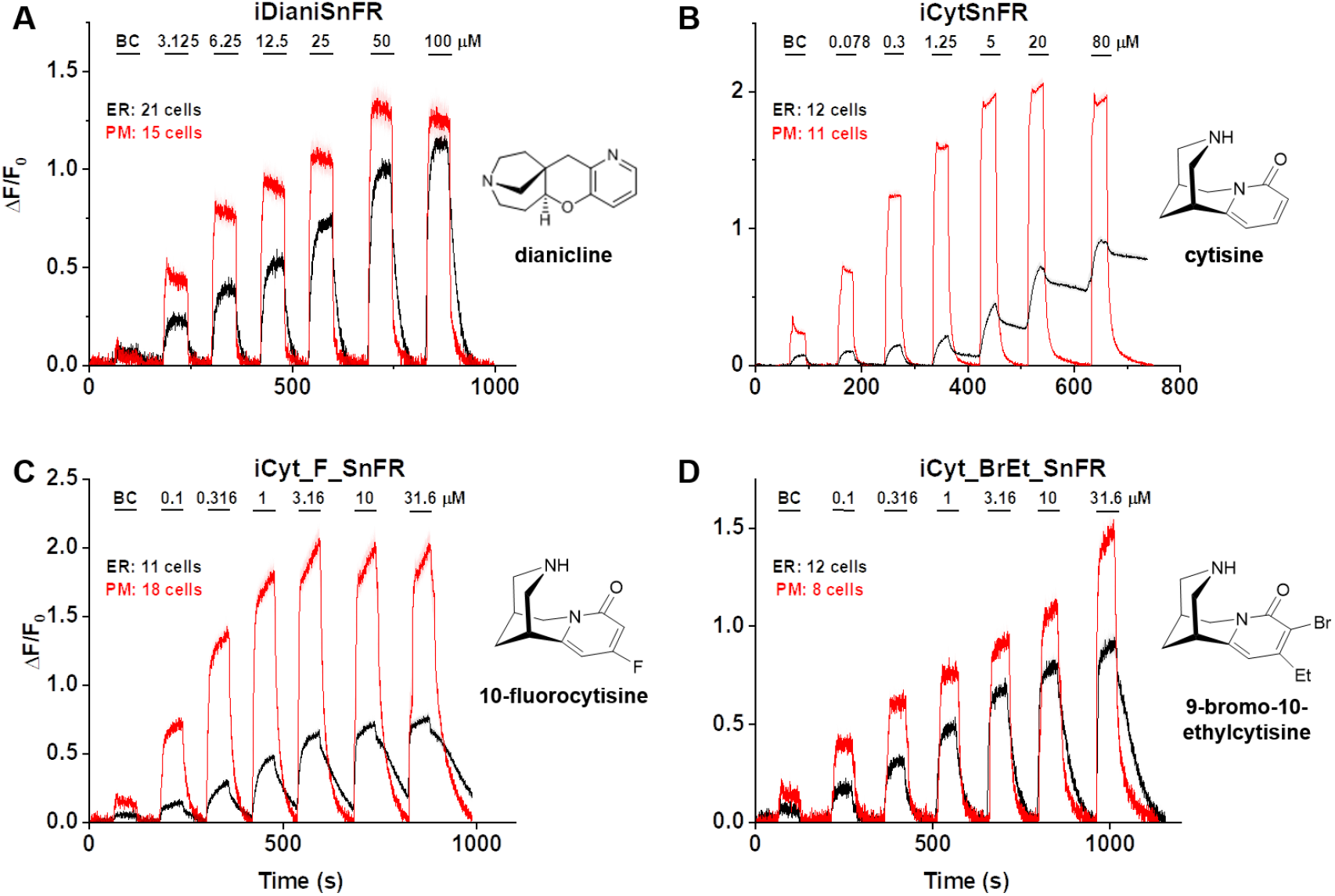
Nicotinic agonist iDrugSnFR dose-response relations in HeLa cells. **(A-D)** Each iDrugSnFR detects its drug partner at the PM and ER of HeLa cells at the concentrations sampled. BC = Buffer control. SEM of data are indicated by semi-transparent shrouds around traces where trace width is exceeded. **(A)** iDianiSnFR detects dianicline with a return to baseline fluorescence between drug applications. **(B)** iCytSnFR detection at the PM returns to baseline fluorescence between applications, while detection at the ER shows incomplete wash-in and washout. **(C)** iCyt_F_SnFR fluorescence response to the presence of 10-fluorocytisine in the ER also shows an incomplete washout between applications. **(D)** iCyt_BrEt_SnFR detects 9-bromo-10-ethylcytisine with wash-in and washout fluorescence similar to the pattern seen in iDianiSnFR.

**Figure 8.**
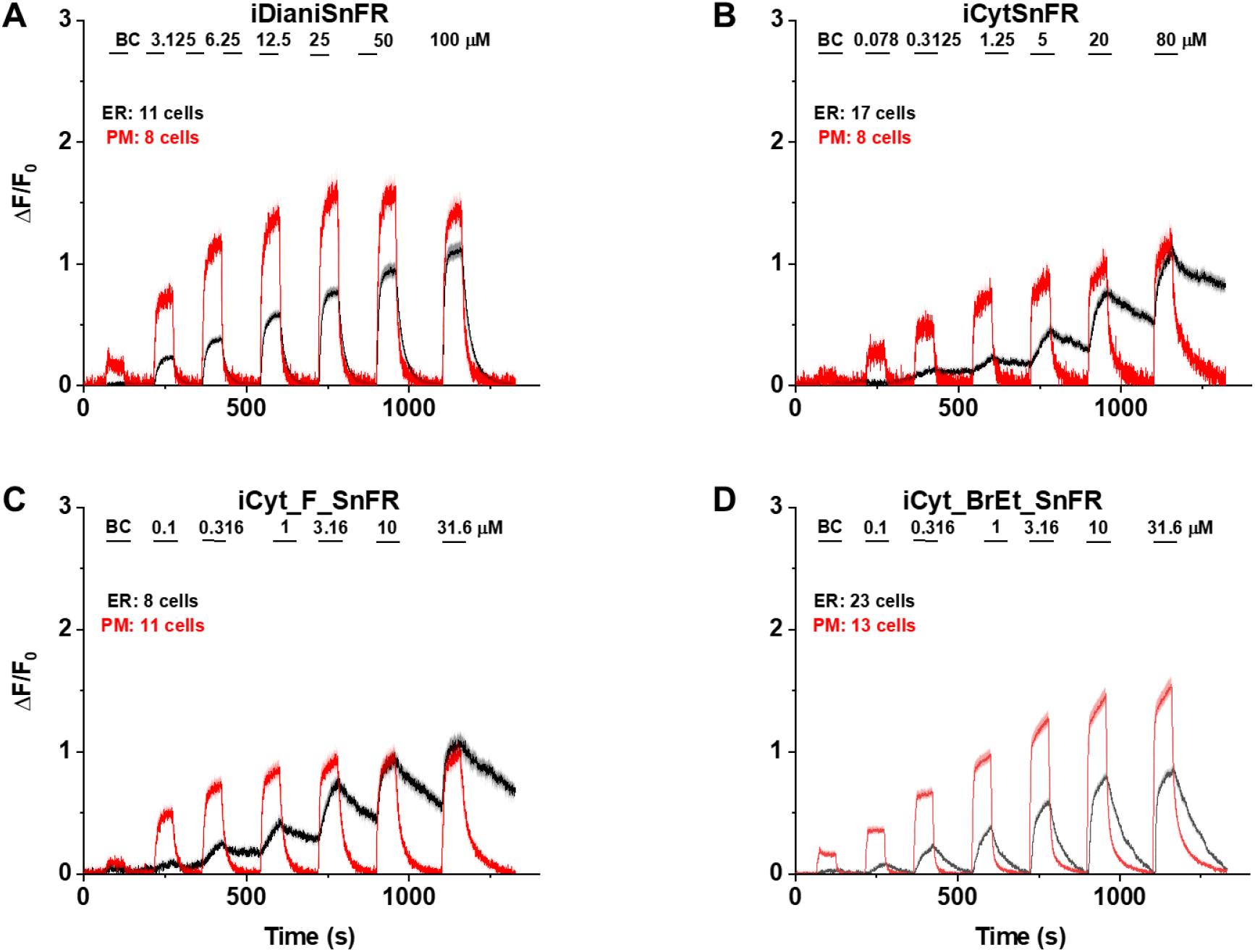
Nicotinic agonist iDrugSnFR dose-response experiments in mouse primary hippocampal neurons transduced with AAV9-hSyn iDrugSnFR. Cultured primary mouse hippocampal neurons were transduced with ER-or PM-targeted constructs. BC = Buffer control. SEM of data are indicated by semi-transparent shrouds around traces where trace width is exceeded. **(A-D)** Each iDrugSnFR detects its drug partner at the PM and ER over the concentrations sampled. **(A**) iDianiSnFR detects dianicline with a return to baseline fluorescence between drug applications. **(B)** iCytSnFR detection at the PM returns to baseline fluorescence between applications, while detection at the ER shows an incomplete washout. **(C)** iCyt_F_SnFR fluorescence response to the presence of 10-fluorocytisine in the ER also shows an incomplete washout between applications. **(D)** iCyt_BrEt_SnFR_ER detects 9-bromo-10-ethylcytisine with a wash-in and decay intermediate between iDianiSnFR and the other two cytisine derivatives.

iDianiSnFR showed a robust response to dianicline at the PM and the ER in HeLa cells across a range of concentrations (3.125-100 µM) and the speed was nearly limited by solution exchanges; there was a clear return to baseline fluorescence upon washout on the order of seconds after each drug application. At 100 µM, the PM and ER have a ΔF/F_0_ of ~1.2, but at lower concentrations, the ER displayed 30–75% of the signal detected at the PM, which may indicate a difference in membrane crossing (Fig. 7A). Imaging in primary mouse hippocampal neurons demonstrated a similar trend (Fig. 8A).

Cytisine showed slower entry into and exit from the ER of HeLa cells. The iCytSnFR_PM construct detected cytisine at concentrations from 0.078–80 µM and demonstrated a return to baseline fluorescence upon washout on the order of seconds after each drug application, reaching a maximum ΔF/F_0_ of ~2 at concentrations above 5 µM (Fig. 7B). In contrast to the _PM construct, the iCytSnFR_ER construct only detected cytisine with a ΔF/F_0_ above the buffer control in the range of concentrations from 1.25–80 µM with a ΔF/F_0_ which was 25–50% of the maximum ΔF/F_0_ detected at the PM. Additionally, in the range of detectable concentrations of cytisine, the washout of cytisine was much slower than solution changes (Fig. 7B). The incomplete washout persists even after several minutes and corresponds with previous suggestions that cytisine has low membrane permeability, as evidenced by its low brain penetration (Rollema, et al., 2010).

In primary mouse hippocampal neurons, iCytSnFR detection of cytisine exhibited the same kinetic trends seen in HeLa cell experiments (Fig. 8B). During cytisine application (60 s) from 0.078-80 µM, the iCytSnFR_PM fluorescence nearly reached a plateau, and during the washout (90-180 s), the fluorescence decayed back to baseline, though the decay slowed after removal of higher [cytisine]. The _PM construct reached a maximum ΔF/F_0_ of ~1.25 at 80 µM, which was approximately 60% of the signal observed in HeLa cell experiments (Fig. 8B). The iCytSnFR_ER detection of cytisine in the ER reflected the trends seen in HeLa cells, with incomplete cytisine wash-in phases and prolonged cytisine washout phases. One observable difference was that the maximum ΔF/F_0_ (~1.25) of iCytSnFR_ER reached a similar maximum to that of iCytSnFR_PM in neurons, which was not observed in HeLa cell experiments (Fig 7B).

In preliminary HeLa cell experiments with varenicline applied to iCytSnFR, we found much slower kinetics that differed little between the _ER and _PM constructs (data not shown). These findings, which vitiated the use of the iCytSnFR:varencline pair in the cellular experiments, are consistent with the markedly slow kinetics of varenicline-iCytSnFR interactions in the microperfusion experiments (see above).

iCyt_F_SnFR targeted to the PM and ER showed characteristics similar to iCytSnFR in HeLa cells. The _PM construct detected 10-fluorocytisine across a range of concentrations with a return to baseline fluorescence between applications, while the _ER construct detected 10-fluorocytisine with ΔF/F_0_ values that were only 25–33% of those detected at the PM (Fig. 7C). Similar to the iCytSnFR_ER detection of cytisine, the iCyt_F_SnFR_ER detection of 10-fluorocytisine was much slower than solution changes and did not return to baseline between applications, though the washout occurs on the order of minutes, rather than tens of minutes as with iCytSnFR_ER (Fig. 7C). The difference in PM and ER detection of 10-fluorocytisine again shows decreased membrane permeability into HeLa cells compared to other drugs we have examined with other iDrugSnFRs. Overall, the detection of 10-fluorocytisine with iCyt_F_SnFR in primary hippocampal culture resembled our data with iCyt_F_SnFR in HeLa cells. Nevertheless, there were distinct differences (Fig. 8C), such as a decreased maximum ΔF/F_0_ in the iCyt_F_SnFR_PM construct and a similar maximum ΔF/F_0_ of ~1 for both the _ER and _PM constructs. Additionally, the decay of the iCyt_F_SnFR responses lasted tens of minutes, resembling the iCytSnFR_ER data in primary hippocampal culture.

9-bromo-10-ethylcytisine showed a kinetic profile resembling dianicline. iCyt_BrEt_SnFR_PM responses to 9-bromo-10-ethylcytisine (0.1-31.6 µM) were nearly limited by solution exchanges with a return to baseline fluorescence on the order of seconds, and a maximum ΔF/F_0_ of ~1.5 at 31.6 µM. iCyt_BrEt_SnFR_ER also detected 9-bromo-10-ethylcytisine over this range of concentrations and returned to baseline fluorescence between applications (Fig. 7D). ΔF/F_0_ values for iCyt_BrEt_SnFR_ER were 50–75% of the ΔF/F_0_ values detected by iCyt_BrEt_SnFR_PM, which indicated that 9-bromo-10-ethylcytisine crossed into and out of cells readily (Fig. 7D). Imaging in primary mouse hippocampal neurons revealed the same trend (Fig. 8D).

To more fully examine the membrane-crossing properties of the nicotinic agonists, we recorded the fluorescence waveforms for several drugs at concentrations between 0.1-3.16 µM with much longer application times and washout times than in the above experiments (Supplementary Fig. 5). With these conditions, the fluorescence signals suggested complete washout of each nicotinic agonist from the ER of HeLa cells. However, it is noteworthy that even when applied at concentrations as low as 0.1 µM and 0.316 µM, cytisine and 10-fluorocytisine require washout times of several min from the ER. In contrast, the iDrugSnFR localized to the PM shows a rapid return to baseline after drug application.

Because the data of Fig. 7B and Fig. 8B indicated that iCytSnFR_PM functions as expected from stopped-flow and millisecond perfusion, we applied additional experiments to ensure that our observations of drug entry and exit from the ER were not the result of idiosyncratic biosensor function or folding in the ER. iCytSnFR and iCyt_F_SnFR both bind nicotine in the same concentration range as cytisine (though with lower ΔF/F_0_). After transfection of _PM and _ER constructs for each sensor into HeLa cells, we performed time-resolved imaging for pulses of 0.1-31.6 µM nicotine (Fig. 7-figure supplement 2). These nicotine waveforms resembled those already published with iNicSnFR3a and iNicSnFR3b (Shivange, et al., 2019), confirming that iCytSnFR_ER functions as expected with a more permeant nicotinic drug. Thus, the slower kinetics for iCytSnFR_ER with-cytisine and iCyt_F_SnFR_ER with 10-fluorocytisine arise because these drugs cross membranes more slowly.

To examine localization of the _PM and _ER constructs at higher optical resolution, we imaged HeLa cells and primary mouse hippocampal culture using a spinning disk laser scanning inverted confocal microscope. As previously observed, ER-targeted iDrugSnFR was retained in the ER (Fig. 7-figure supplement 1, Fig. 8-figure supplement 1) (Shivange, et al., 2019). IDrugSnFR targeted to the PM showed correct localization, with some iDrugSnFR observed in the cell interior (most likely as part of the cellular membrane trafficking system (Fig. 7-figure supplement 1; Fig. 8-figure supplement 1).

Several complexities in the HeLa cell and neuron experiments imposed uncertainties on our kinetic analyses. These complexities include the limitations of solution changes, diffusion within cytoplasm, unknown mixing at the surface facing the coverslip, and corrections for baseline drift due to bleaching. We restrict the quantitative comparisons to the estimate that cytisine and 10-fluorocytisine cross the membrane > 30 fold more slowly than the other drugs tested.

## Discussion

### Membrane permeation of molecules with low logD_pH7.4_

The experiments show, to our knowledge, the first time-resolved measurements of membrane permeation for drugs in the logD_pH7.4_ range less than -1. Most orally available drugs have logD_pH7.4_ values between 2 and 4 (Smith, Allerton, Kalgutkar, van de Waterbeemd, & Walker, 2012). Cytisine, varenicline, dianicline, and the cytisine analogs studied here have calculated membrane partition coefficients some 3 to 6 orders of magnitude lower. These values and their order vary according to the algorithm, partially because of uncertainties in predicting pK_a_ (Pienko, et al., 2016); here we provide values calculated by Chemicalize (see Methods): 10-fluorocytisine, -2.70; cytisine, -2.64; dianicline, -1.29; varenicline, -1.27, 9-bromo-10-ethylcytisine, -1.13; It is remarkable that drugs with such low calculated partition coefficients do cross membranes on a time scale of seconds (9-bromo-10-ethylcytisine, varenicline, dianicline) to minutes (10-fluorocytisine, cytisine). According to some (but not all) algorithms, the calculated logD_pH7.4_ values fall in the same two classes as the measured kinetics of membrane permeability: 10-fluorocytisine and cytisine are the slowest, and only these two agonists have logD_pH7.4_ values < -2. These observations support previous work suggesting that differences among chemical properties of nicotinic partial agonists correlate with drug permeation into the cerebrospinal fluid (CSF) after peripheral administration in mice (Rollema, et al., 2010).

### The iDrugSnFR paradigm

The iDrugSnFRs are sensitive enough to allow experiments near the experimentally determined (or otherwise projected) concentration in the human blood and CSF (Astroug, Simeonova, Kassabova, Danchev, & Svinarov, 2010; Jeong, Sheridan, Newcombe, & Tingle, 2018; Rollema, et al., 2010). The iDrugSnFRs have the advantage that they measure free aqueous ligand concentration (“activity”), as sensed by nAChRs. Targeting sequences provide for visualization within the lumen of organelles—here, the ER.

The experiments do not use radiolabeled drugs, *in vivo* microdialysis or other experiments on live animals, or mass spectrometry-liquid chromatography instruments. Once protein design has given an entry into a class of iDrugSnFRs, straightforward optimization at the binding site produces the desired, selective iDrugSnFRs for individual molecules. For drugs that bind at orthosteric cholinergic sites (both nicotinic and muscarinic), we anticipate that a collection of tens, rather than hundreds, of iDrugSnFRs will suffice to detect all present and future ligands. The experiments use standard, modest-power fluorescence microscopes. Cultured cell lines yield data comparable to cultured neurons.

We comment on varenicline. None of the biosensors in Table 1 were evolved to bind varenicline; yet it binds to some iDrugSnFRs with nanomolar EC_50_. Only iDianiSnFR, which lacks His68, binds varenicline with EC_50_ > 10 μM. Even tighter binding has been achieved with varenicline derivatives at mutated ligand-gated channels (Magnus, et al., 2019). On the one hand, the cellular experiments described here cannot use iDrugSnFR pairs with dissociation rate constants less than ~ 0.1 s^-1^, corresponding to an EC_50_ of less than ~ 100 nM. On the other hand, all known neural drugs leave the human body and brain much more slowly, with rates determined primarily by metabolism; even “fast” nicotine metabolizers display time constants of ~ 1200 s (Dempsey, et al.). Highly sensitive, tightly binding, reagentless iDrugSnFRs will be used in studies on personal pharmacokinetics in biofluids.

### Structure-function relations for nicotinic and other iDrugSnFRs

This study shows that the amine group of nicotinic ligands makes equidistant cation-π interactions with two tyrosine residues (Tyr65, Tyr357), and this is confirmed by higher-resolution (1.5 to 1.7 Å) structures of varenicline, acetylcholine, and choline crystallized with isolated PBP moieties (PDB 7S7X, S6V1R, 7S7Z, respectively; see also 3R6U, 6EYQ, and 3PPQ). Cation-π interactions also occur for cholinergic and/or nicotinic ligands in nAChRs (Morales-Perez, Noviello, & Hibbs, 2016; Post, Tender, Lester, & Dougherty, 2017), the acetylcholine-binding protein (Celie, et al., 2004), PBPs (Schiefner, et al., 2004), and muscarinic receptors (Haga, et al., 2012). We also observe that the protonated amine of varenicline makes a hydrogen bond to a backbone carbonyl group, another similar theme in acetylcholine binding protein (Celie, et al., 2004) and nAChRs (Xiu, Puskar, Shanata, Lester, & Dougherty, 2009).

This study presents a general step forward in understanding the structure-function relations of iDrugSnFRs. The chromophore in the cpGFP moiety of most present iDrugSnFRs (this paper, iGluSnFR, iSeroSnFR) contains a tyrosine in an extended π system (Ormo, et al., 1996; Tsien, 1998). The photophysics of the chromophore depends strongly on the surrounding water molecules and side chains (Brejc, et al., 1997; Tsien, 1998). We found that Glu78 in Linker 1 changes its orientation: in the liganded state, it interacts with two positively charged residues (Lys97 and Arg99) on the surface of the cpGFP; and in the apo state, Glu78 has moved ~ 14 Å to form a hydrogen bonding interaction with the tyrosine moiety of the chromophore. Presumably the liganded state of iNicSnFR3adt allows for a water molecule to hydrogen bond with the hydroxy group of the chromophore, promoting its fluorescence; but this water molecule is replaced by protonated Glu78 in the unliganded state, which leads to nonfluorescent state of cpGFP, as suggested by (Nasu, et al., 2021).

While we cannot resolve the protonation-deprotonation event, the available functional data show good support for its occurrence, as follows. (1) The apo form of the iDrugSnFR increases its F_0_ by ten-fold per pH unit (Shivange, et al., 2019), as though when deprotonated, Glu78 leaves the “candle snuffer” position and moves to make the salt bridges with Lys97 and Arg99. (2) The EC_50_ for the ligand decreases by ten-fold per pH unit (Shivange, et al., 2019), as though the conformation of the linker that forms the salt-bridge form is also the closed, liganded, fluorescent form of the PBP. Other observations favor the crucial role of the Glu78-chromophore interaction. (3) Only glutamate functions in position 78 of iSeroSnFR (Unger, et al., 2020). (4) The mTurquoise variant in iGluSnFR, which has a tryptophan chromophore, requires entirely different linkers (Marvin, et al., 2018).

### Challenges at the intersection of pharmaceutical science and nicotine addiction science

Our measurements show that nicotinic ligands with logD_pH7.4_ < ~ -2 cross membranes much more slowly than do ligands with logD_pH7.4_ > ~ -2. These measurements have two, possibly opposing, implications for future smoking cessation drugs. On the one hand, α4β2 agonists that enter the ER, like nicotine and varenicline, upregulate nAChRs (Turner, Castellano, & Blendy, 2011), which may be necessary and sufficient for addiction (Henderson & Lester, 2015); and maintenance of upregulation by varenicline may help to explain its suboptimal quit rate. On the other hand, ligands that do not enter the ER are also unlikely to enter the brain and therefore unlikely to be useful for smoking cessation.

Smoking cessation drugs must also contend with other ER-based processes. (1) Most drug metabolism takes place in the ER; and (2) upregulation occurs at a sustained agonist concentration in the ER some hundredfold lower than the extracellular concentrations that transiently activate α4β2 nAChRs (Kuryatov, Luo, Cooper, & Lindstrom, 2005).

Given these challenges, further progress may be possible now that we have two types of real-time, living cellular preparations. (1) For decades, cellular preparations have been available to measure nAChR pharmacodynamics and upregulation. (2) Now, the iDrugSnFRs present a paradigm to measure cellular and subcellular pharmacokinetics. The iDrugSnFR paradigm will be useful beyond the explicit case of nicotine addiction, with application to other exogenous neural drugs.

## Methods

### Crystallography

The gene encoding the full-length biosensor iNicSnFR3a was previously cloned into a bacterial expression vector (Shivange, et al., 2019). To improve crystallization, we deleted the N-terminal HA tag and the N-terminal Myc tag, forming the constructs with the suffix “dt”. These deletions were carried out with the Q5 Site-Directed Mutagenesis Kit (New England Biolabs, Ipswich, MA). All proteins were overexpressed in *E. coli* BL21-gold (DE3) cells (Agilent Technologies, Santa Clara, CA) using ZYM-5052 autoinduction media (Studier, 2005). Cells were collected by centrifugation and stored at -80 °C until use.

For purification, frozen cell pellets were resuspended in lysis buffer containing 100 mM NaCl, 20 mM Tris, pH 7.5, 20 mM imidazole, pH 7.5, 5 mM β-mercaptoethanol (BME), lysozyme, DNase, and protease inhibitor tablet. The resuspended cells were lysed by freezing and thawing using liquid nitrogen and a room temperature water bath for 3 cycles. Intact cells and cell debris were removed by centrifugation at ~20,000x g for 40 min at 4 °C. The supernatant was collected and loaded onto a prewashed Ni NTA column with wash buffer at 4 °C. Ni NTA wash buffer contained 100 mM NaCl, 20 mM Tris, pH 7.5, 30 mM imidazole, pH 7.5, and 5 mM BME. Elution was achieved using the same buffer with 300 mM imidazole, pH 7.5. The eluted sample was further purified by size exclusion chromatography using HiLoad 16/60 Superdex 200 in the same buffer without imidazole and BME. Peak fractions were collected and concentrated to ~50 mg/ml with Amicon Ultra 15 filter unit (Millipore, Burlington, MA) with 10kDa cutoff.

For all constructs, initial crystallization screening was carried out with 40 mg/ml protein in the presence and absence of 10 mM nicotine or varenicline. iNicSnFR3adt crystallized separately with 10 mM nicotine and varenicline in PACT premier (Molecular Dimensions, Sheffield, England), condition #96 with 0.2 M sodium malonate dibasic monohydrate, 0.1 M Bis-Tris Propane, pH 8.5, and 20% polyethylene glycol (PEG) 3,350 at 20 °C. Crystals of iNicSnFR3adt grew within two weeks of crystallization in a hexagonal rod shape with dimensions of ~ 80 μm x 80 μm x 300 μm. Crystals were harvested and cryo-protected in 25% ethylene glycol, 0.2 M sodium malonate dibasic monohydrate, 0.1 M BisTrisPropane pH 8.5, and 20% PEG 3350. Phase information was obtained through soaking with KI before cryo-protection. The unliganded iNicSnFR3adt crystallized in Morpheus (Molecular Dimensions), condition #92 with 2.5% PEG 1,000, 12.5% PEG 3,350, 12.5% 2-methyl-2,4-pentanediol, 0.02 M of each amino acid, and 0.1 M MOPS/HEPES-Na, pH 7.5 at 23 °C with no further optimization.

X-ray datasets were collected at Stanford Synchrotron Radiation Laboratory beamline 12-2 and Advanced Light Source beamline 5.0.2 using Pilatus 6M detectors. All datasets were processed and integrated with XDS (Kabsch, 2010) and scaled with Aimless (Winn, et al., 2011). For iNicSnFR3adt, molecular replacement was carried out using domains of the unliganded structure (PDB ID: 6EFR) with Phaser in Phenix (Adams, et al., 2010). The experimental phase information of KI-soaked crystals of iNicSnFR3adt was obtained with MR-SAD using AutoSol in Phenix (Adams, et al., 2010). Molecular replacements of the remaining structures were carried out with the refined model of iNicSnFR3adt. Iterative refinement and model building cycles for all structures were carried out separately with phenix.refine in Phenix (Adams, et al., 2010) and Coot (Emsley, Lohkamp, Scott, & Cowtan, 2010).

### Directed evolution of iDrugSnFR proteins using bacterial-expressed protein assays

Starting with iAChSnFR and intermediate biosensor constructs of that sensor, we constructed and optimized iDrugSnFRs for each drug partner during iterative rounds of SSM as previously described (Bera, et al., 2019; Shivange, et al., 2019). We utilized the 22-codon procedure including a mixture of three primers, creating 22 unique codons encoding the 20 canonical amino acids (Kille, et al., 2013). The 22-codon procedure yields an estimated > 96% residue coverage for a collection of 96 randomly chosen clones.

A Tecan Spark M10 96-well fluorescence plate reader (Tecan, Männedorf, Switzerland) was used to measure baseline and drug-induced fluorescence (F_0_ and ΔF, respectively). Bacterial lysates were tested with excitation at 485 nm and emission at 535 nm. Lysates were also measured against choline to evaluate potential endogenous intracellular binding. Promising clones were amplified and sequenced. The optimally responding construct in each round of SSM was used as a template for the next round of SSM.

S-slope allows for comparison between iDrugSnFRs with differing *ΔF*_*max*_*/F*_*0*_ values (Bera, et al., 2019) at the beginning of the dose-response relation, which is usually the pharmacologically relevant range. With lysates or purified protein, which allow complete dose-response relations, the Hill coefficient is near 1.0. We therefore calculated

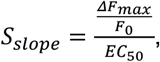

in units of µM^-1^.

### Measurements on purified iDrugSnFRs

Biosensors selected for further study were purified using a His_6_ sequence using an ÄKTA Start FPLC (GE Healthcare, Chicago, IL) as previously described (Shivange, et al., 2019). Performance of protein quantification and dose-response relations for drug-sensor partners was also as previously described (Shivange, et al., 2019). Where appropriate, we corrected for depletion of the ligand by binding with the equation,

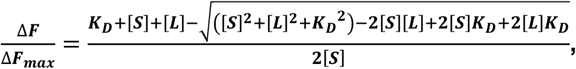

where *K*_*D*_ is the ligand-sensor equilibrium dissociation constant (we assume that *K*_*D*_ = *EC*_*50*_), *[S]* is the iDrugSnFR protein concentration (typically 100 nM), and *[L]* is the nominal ligand concentration.

### Isothermal titration calorimetry

Experiments were performed on an Affinity ITC (TA instruments, New Castle, DE) at 25 °C. The iDrugSnFR protein was buffer-exchanged into 3x PBS, pH 7.0. The nicotinic agonists were dissolved in the same buffer. 800 µM cytisine (Sigma Aldrich, Munich, Germany) was titrated into 80 µM iCytSnFR, 160 µM 10-fluorocytisine was titrated into 16 µM iCyt_F_SnFR. 470 µM 9-bromo-10-ethylcytisine was titrated into 47 µM iCyt_BrEt_SnFR. 1.5 mM dianicline (Tocris, Bio-Techne, Minneapolis, MN) was titrated into 150 µM iDianiSnFR. Analysis, including correction for changes in enthalpy generated from the dilution of the ligands, was performed using a single-site binding model in the manufacturer’s Nanoanalyze software.

### Stopped-flow kinetic analysis

Kinetics were determined by mixing equal volumes of 0.2 µM iDrugSnFR protein (in 3x PBS, pH 7.0) with varying concentrations of cognate ligand in an Applied Photophysics (Surrey, United Kingdom) SX20 stopped-flow fluorimeter with 490 nm LED excitation and 510 nm long-pass filter at room temperature (22 °C). “Mixing shots” were repeated five times and averaged (except for 100 s experiments, which were collected only once). Standard deviations are not included on the plots, but are nearly the same size as the data markers. The first 3 ms of data were ignored because of mixing artifacts and account for the dead time of the instrument.

Data were plotted and time courses were fitted, when possible, to a single exponential, and k_obs_ was plotted as a function of [ligand]. The linear portion of that graph was fit, with the slope reporting k_1_ and the y-intercept reporting k_-1_. When the time course did not fit well to a single rising exponential, it was fitted to the sum of two increasing exponentials, and the first rise (k_obs1_) was treated as above to determine k_1_ and k_-1_.

### Expression in mammalian cells

We constructed two variants of each iDrugSnFR for expression in mammalian cells. The plasma membrane (suffix _PM) and endoplasmic reticulum (suffix _ER) variants were constructed by circular polymerase extension cloning (Quan & Tian, 2009). To create the _PM constructs, we cloned the bacterial constructs into pCMV(MinDis), a variant of pDisplay (ThermoFisher Scientific, Waltham, MA) lacking the hemagglutinin tag (Marvin, et al., 2013). To generate the _ER constructs, we replaced the 14 C-terminal amino acids (QVDEQKLISEEDLN, including the Myc tag) with an ER retention motif, QTAEKDEL (Shivange, et al., 2019).

We transfected the iDrugSnFR cDNA constructs into HeLa and HEK293T cells. Cell lines were purchased from ATCC (Manassas, VA) and cultured according to ATCC protocols. We purchased new aliquots of the cell lines listed above at six-month intervals to ensure reproducibility. Mycoplasma contamination was assayed at six-month intervals and was negative over the course of these experiments. For chemical transfection, we utilized either Lipofectamine 2000 or Lipofectamine 3000 (ThermoFisher Scientific), following the manufacturer’s recommended protocol. Cells were incubated in the transfection medium for 24 h and imaged 24 – 48 h after transfection.

### Millisecond timescale microperfusion

HEK293T cells were imaged using a Nikon (Tokyo, Japan) DIAPHOT 300 with a Zeiss 63X objective (1.5 NA). Because the ligand concentration after micro-iontophoretic drug application (Shivange, et al., 2019) is unknown, we applied drugs with a laminar-flow microperfusion (Model SS-77B Fast-Step perfusion system (Warner Instruments, Holliston, MA). In an array of three square glass capillaries (600 µ i.d.), the center capillary contained vehicle (Hanks buffered salt solution, HBSS) plus drug, while the two outer capillaries contained vehicle only. Vehicle also flowed from a separate input connected to the bath perfusion system. Solution exchange, measured by loading the center capillary with dye, had a time constant of 90 ± 20 ms (n = six trials).

We used Fiji ImageJ and Origin Pro 2018 (OriginLab, Northampton, MA) to fit the rise and decay of the iCytSnFR_PM drug response to the sum of one or two exponential components. An F-test determined whether two exponential components fit the data significantly better than one (p < 0.05). Statistical comparisons between groups were carried out using ANOVA.

### AAV production and transduction in primary mouse hippocampal neuronal culture

The adeno-associated virus plasmid vector AAV9-hSyn was described previously (Challis, et al., 2019). Virus was purified using the AAVpro Purification Kit (TakaraBio USA). Mouse embryo dissection and culture were previously described (Shivange, et al., 2019). About 4 days after dissection, we transduced the _ER construct at an MOI of 0.5 to 5 × 10^4^; and separately, the _PM construct was transduced at an MOI of 0.5 to 1 × 10^5^. Neurons were imaged ~2-3 weeks post-transduction.

### Time-resolved fluorescence measurements in live mammalian cells and primary mouse hippocampal neuronal culture

Time-resolved dose-response imaging was performed on a modified Olympus IX-81 microscope (Olympus microscopes, Tokyo, Japan), in widefield epifluorescence mode using a 40X lens. Images were acquired at 2 – 4 frames/s with a back-illuminated EMCCD camera (iXon DU-897, Andor Technology USA, South Windsor, CT), controlled by Andor IQ3 software. Fluorescence measurements at λ_ex_ = 470 nm and the epifluorescence cube were as previously described (Shivange, et al., 2019; Srinivasan, et al., 2011).

Solutions were delivered from elevated reservoirs by gravity flow, via solenoid valves (Automate Scientific, Berkeley, CA), then through tubing fed into a manifold, at a rate of 1-2 ml/min. The vehicle was HBSS. Other details have been described (Shivange, et al., 2019; Srinivasan, et al., 2011). Data analysis procedures included subtraction of “blank” (extracellular) areas and corrections for baseline drifts using Origin Pro 2018.

### Spinning disk confocal fluorescence images

HeLa cells and mouse primary hippocampal culture were transfected or transduced as described above. Live-cell images were collected using a Nikon Ti-E spinning disk laser scanning confocal inverted microscope equipped with 100X objective, 1.49 NA (oil), 120 μm WD. The laser wavelength was 488 nm at 15% power. Dishes were imaged in a custom incubator (Okolab, Ottaviano, Italy) at 37° C and 5% CO_2_. Initial images were taken in HBSS. To add drug, we doubled the bath volume by adding HBSS containing drug, using a hand-held pipette. The final drug concentrations: dianicline, 15 µM; cytisine, 10 µM; 10-fluorocytisine, 10 µM; 9-bromo-10-ethylcytisine, 7.5 µM.

### LogD calculations

We used Chemicalize (https://chemaxon.com/products/chemicalize). The software uses algorithms to calculate LogP and pK_a_. The software then calculates

LogD_7.4_ = logP -log[1 + 10^7.4 - pKa^].

### Plasmid availability

We will deposit plasmids with the following cDNAs at Addgene:

iDianiSnFR,
iCytSnFR,
iCyt_F_SnFR,
iCyt_BrEt_SnFR

We will deposit the following plasmids at Addgene:

pCMV(MinDis)-iDianiSnFR_PM,
pCMV(MinDis)-iCytSnFR_PM
pCMV(MinDis)-iCyt_F_SnFR_PM
pCMV(MinDis)-iCyt_BrEt_SnFR_PM
pCMV(MinDis)-iDianiSnFR_ER,
pCMV(MinDis)-iCytSnFR_ER,
pCMV(MinDis)-iCyt_F_SnFR_ER,
pCMV(MinDis)-iCyt_BrEt_SnFR_ER
pAAV9-hSyn-iDianiSnFR_PM,
pAAV9-hSyn-iCytSnFR_PM,
pAAV9-hSyn-iCyt_F_SnFR_PM,
pAAV9-hSyn iCyt_BrEt_SnFR_PM,
pAAV9-hSyn-iDianiSnFR_ER,
pAAV9-hSyn-iCytSnFR_ER,
pAAV9-hSyn-iCyt_F_SnFR_ER,
pAAV9-hSyn iCyt_BrEt_SnFR_ER,

## Acknowledgements

We thank Stefan Petrovic for his stewardship of the isothermal titration calorimeter in the Caltech Center for Molecular Medicine, Jens Kaiser for help with structural studies at the Caltech Molecular Observatory, the Gradinaru lab and Caltech CLOVER Center for help with viral vectors, and Andres Collazo and Giada Spigolon at the Caltech Biological Imaging Facility. We thank Zoe Beatty, Kallol Bera, Eve Fine, Shan Huang, Elaine Lin, Stephen Mayo, Lin Tian, and Elizabeth Unger for advice and guidance. We thank Achieve Life Sciences for a gift of cytisine.

## Funding

California Tobacco-Related Disease Research Program (TRDRP) (27FT-0022), Aaron L. Nichols.

California Tobacco-Related Disease Research Program (TRDRP) (27IP-0057), Henry A. Lester.

California Tobacco-Related Disease Research Program (TRDRP) (T29IR0455), Dennis A. Dougherty.

NIH (GM-123582, DA043829), Henry A. Lester.

NIH (DA049140, GM7616), Anand K. Muthusamy.

Howard Hughes Medical Institute (Loren L. Looger, Jonathan S. Marvin, Douglas C. Rees).

UK Engineering and Physical Sciences Research Council (No. EP/N024117/1), Timothy Gallagher.

Leiden University International Studies Fund (LISF L18020-1-45), Laura Luebbert.

---

**Fiigure 1-figure supplement 1.**
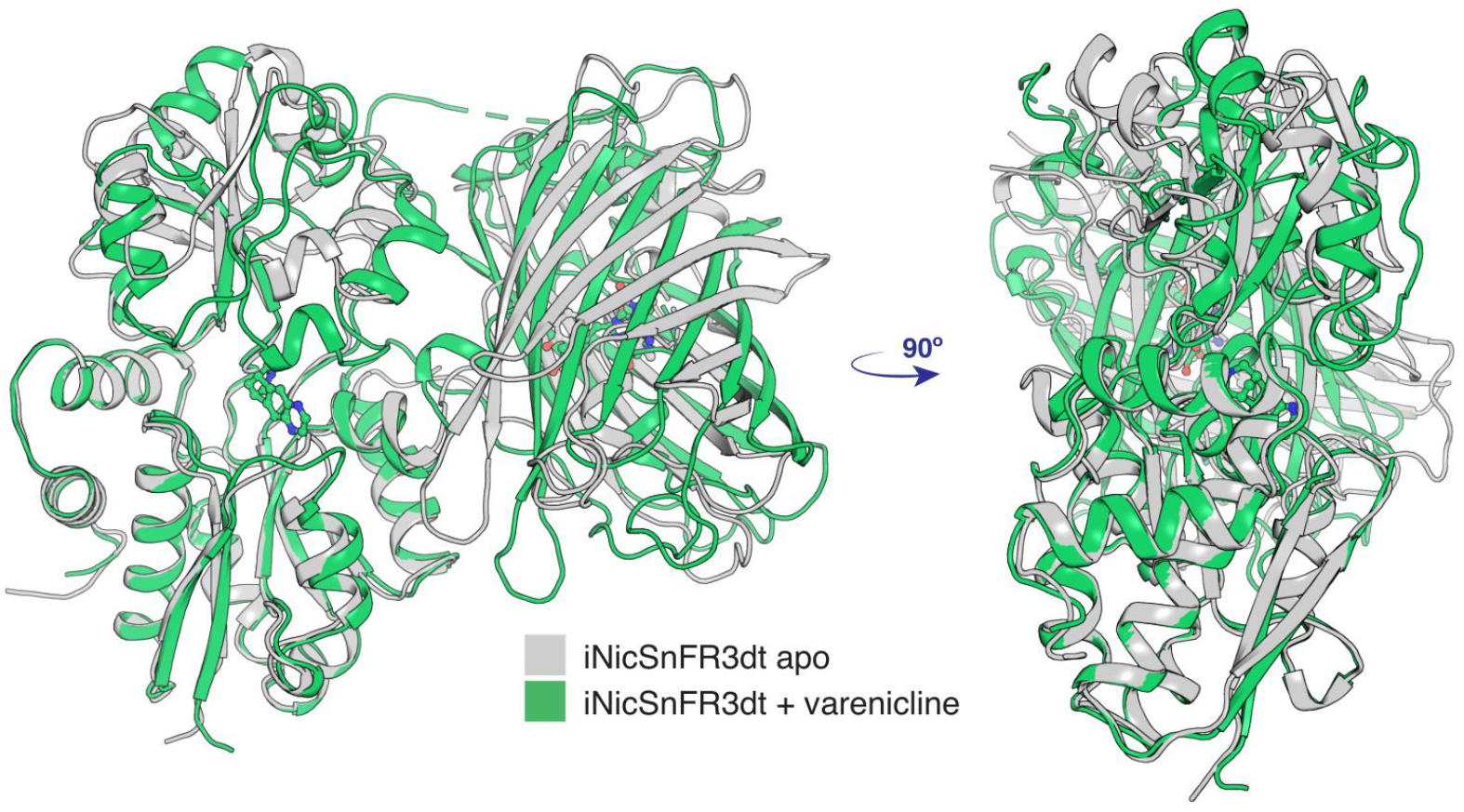
Conformational change of apo (PDB 7S7V) to the liganded, closed form (PDB 7S7T) of iNicSnFR3adt. The bottom lobe of the PBP is superimposed in the two conformations. With respect to the bottom lobe, the “Venus flytrap” conformational change tilts the top lobe of the PBP but does not change its structure (see Supplementary Data). The conformational change also tilts the cpGFP moiety but does not change its structure.

**Fiigure 1-figure supplement 2.**
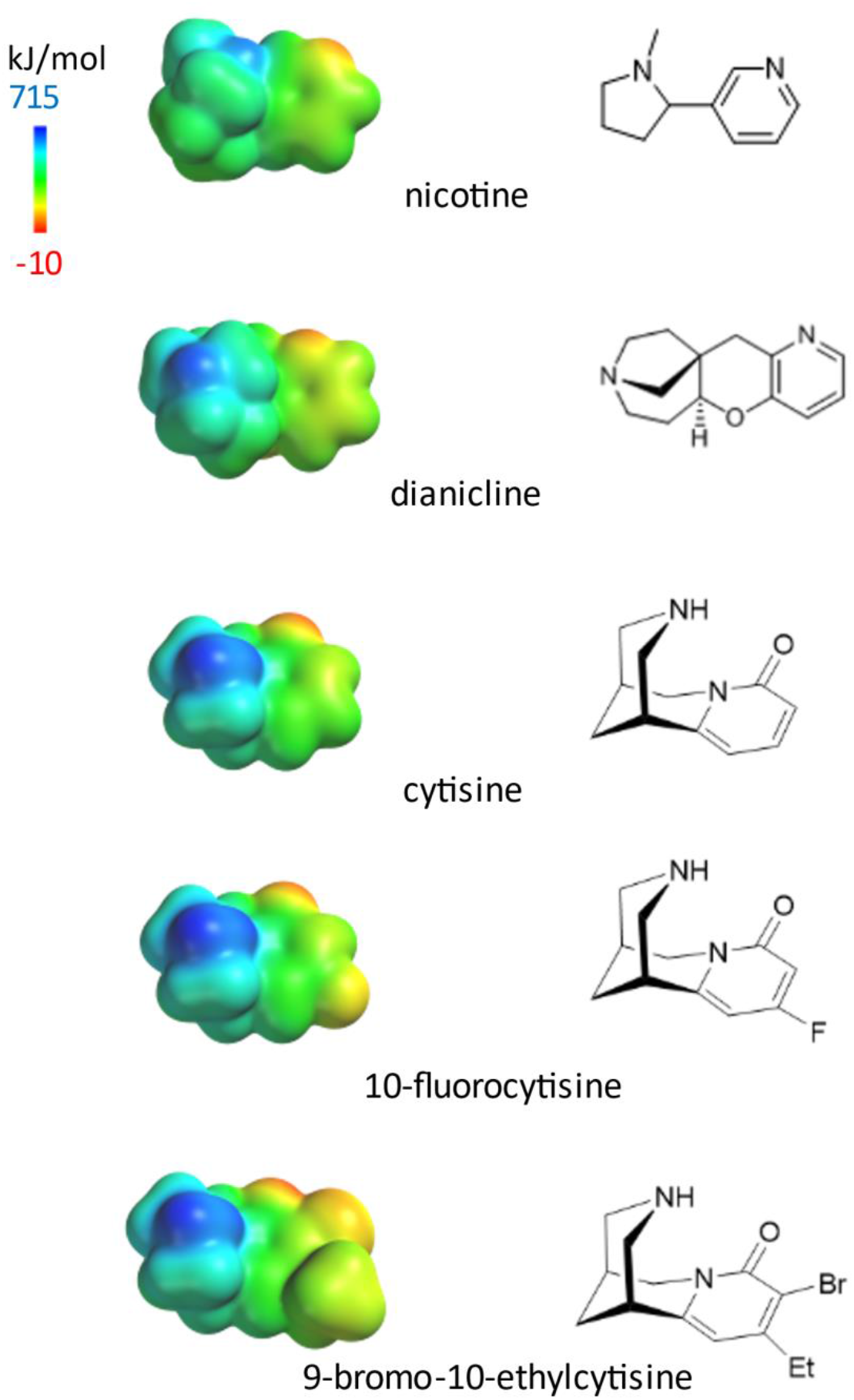
Left, electrostatic surface potential densities for protonated forms of the nicotinic agonists in this study, calculated by SPARTAN at HF/6-31G** theory level. The display ranges from -10 to 715 kJ/mol. The molecules are shown on the same distance scale. Right, bond-line skeletal structures for the deprotonated forms.

**Fiigure 7-figure supplement 1.**
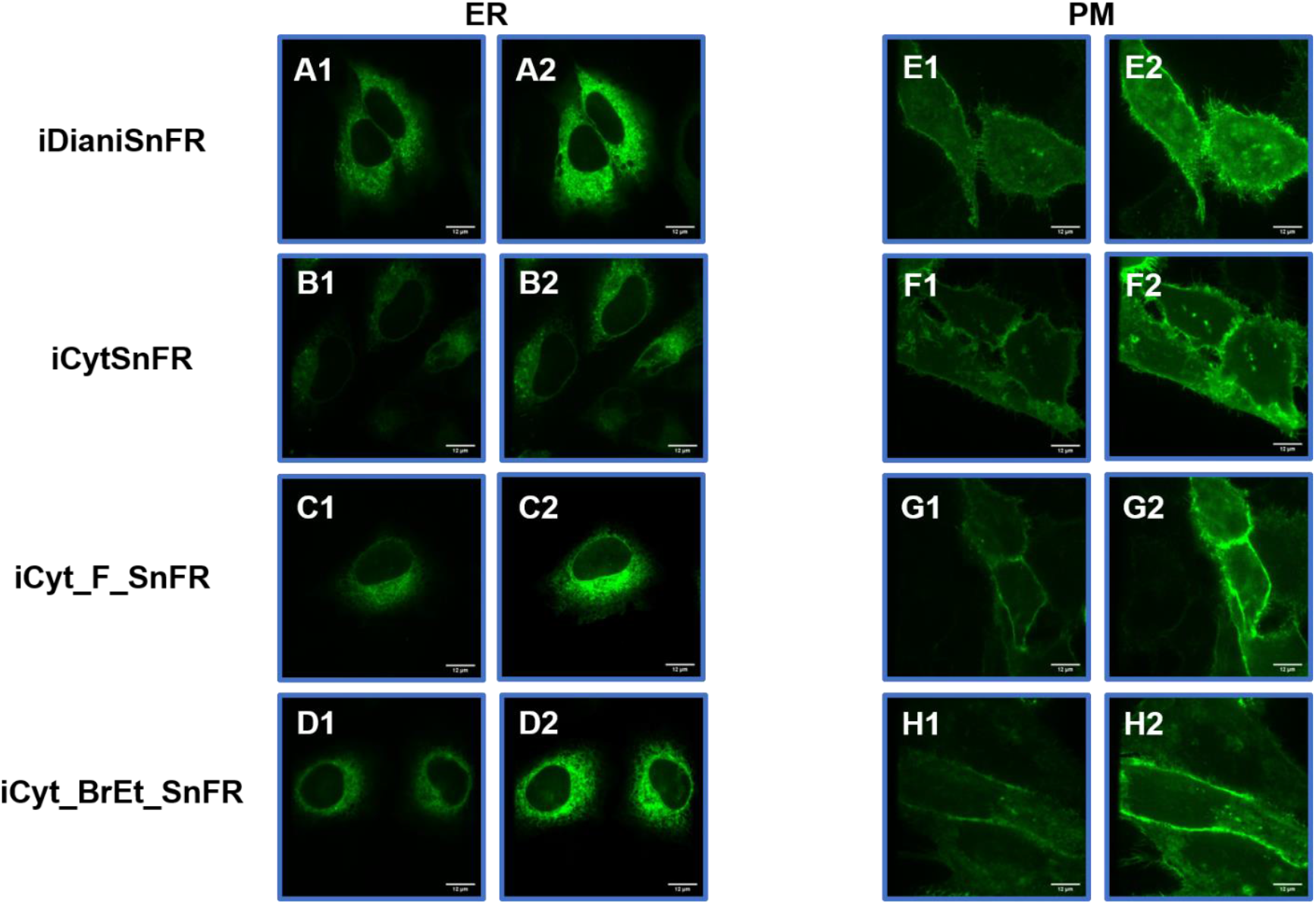
Spinning disk laser scanning confocal inverted microscope images of nicotinic agonist iDrugSnFRs in HeLa cells. ER-targeted constructs of iDianiSnFR, iCytSnFR, iCyt_F_SnFR, iCyt_BrEt_SnFR are shown before **(A1-D1)** and during **(A2-D2)** exposure to each drug partner. ER-targeted iDrugSnFRs show the reticulated ER and dark ovals corresponding to the nucleus. PM-targeted constructs of the same iDrugSnFRs are shown before **(E1-H1)** and after **(E2-H2)** drug introduction. Localization to the PM is robust, with some minimal puncta that may represent inclusion bodies or internal transport.

**Fiigure 7-figure supplement 2.**
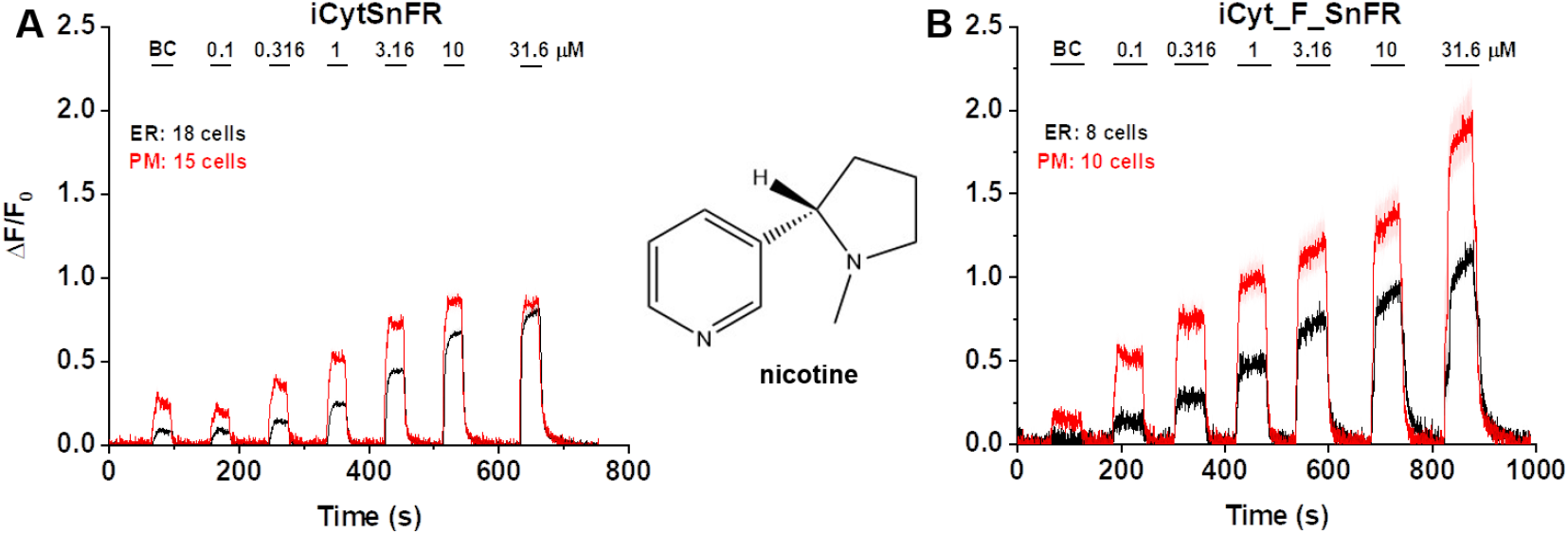
Dose-response relations for iCytSnFR and iCyt_F_SnFR against nicotine in HeLa cells. BC = Buffer control. SEM of data are indicated by semi-transparent shrouds around traces where trace width is exceeded. **(A)** iCytSnFR and **(B)** iCyt_F_SnFR detect nicotine at both the PM and ER. Nicotine enters and exits the ER rapidly over seconds, a direct contrast to the behavior of cytisine and 10-fluorocytisine as detected by their iDrugSnFR partners.

**Fiigure 8-figure supplement 1.**
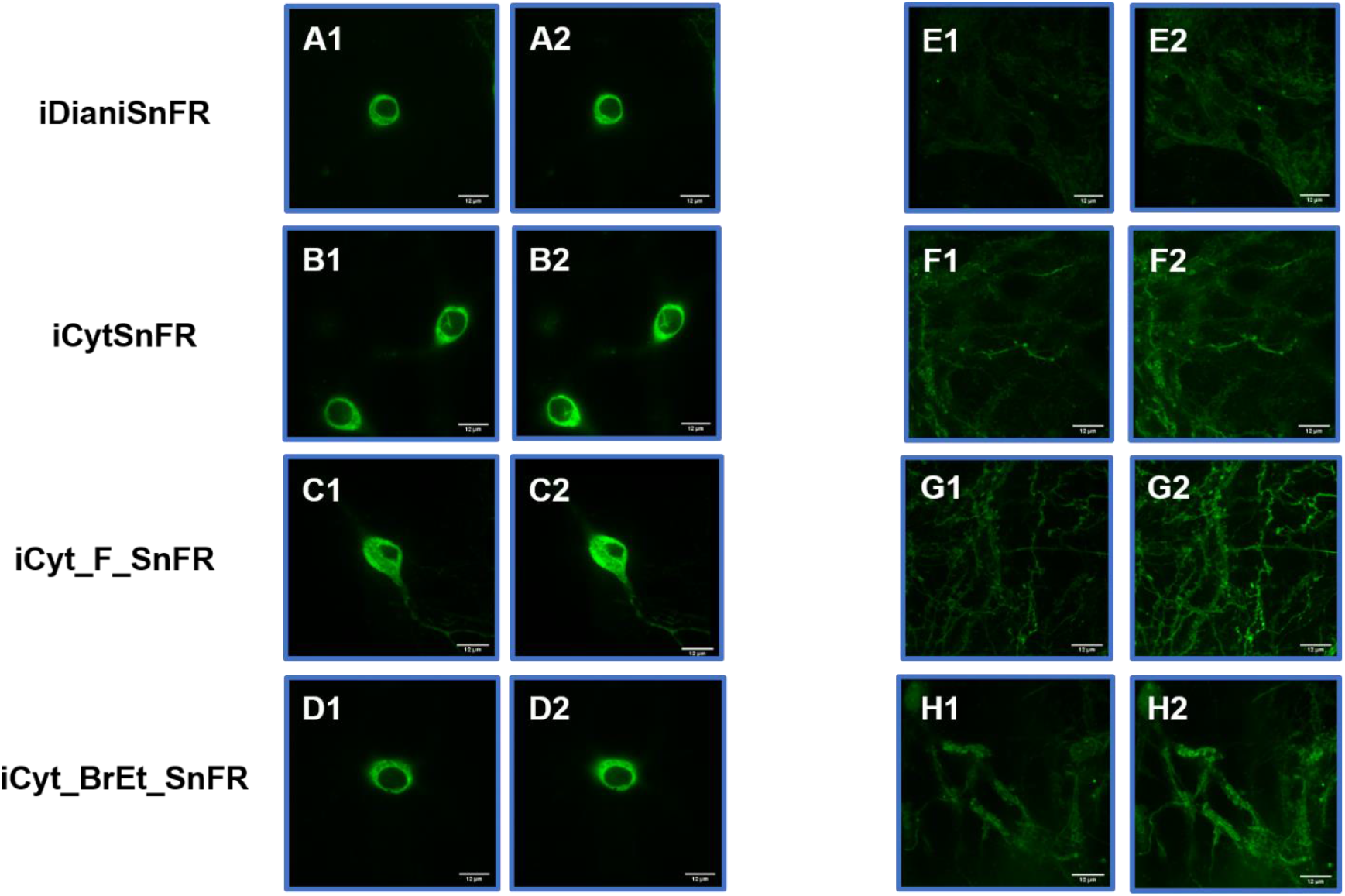
Spinning disk laser scanning confocal inverted microscope images of nicotinic agonist iDrugSnFRs in primary mouse hippocampal neurons. ER-targeted constructs of iDianiSnFR, iCytSnFR, iCyt_F_SnFR, and iCyt_BrEt_SnFR are shown before **(A1-D1)** and during **(A2-D2)** exposure to each drug partner. ER-targeted iDrugSnFRs show the reticulated the ER and dark ovals corresponding to the nucleus. PM-targeted constructs of the same iDrugSnFRs are shown before **(E1-H1)** and after **(E2-H2)** drug introduction. Localization in the PM is robust, with some minimal puncta that may represent inclusion bodies or internal transport.

