## Supplementary Data for "Fluorescence Activation Mechanism and Imaging of Drug Permeation with New Sensors for Smoking-Cessation Ligands"

### iDrugSnFR structural transition

Because all our iDrugSnFRs derive from the *Thermoanaerobacter sp.* X513 choline/betaine binding protein, we asked whether the design process affected conformational changes between the nonfluorescent and fluorescent states. Principal component analysis showed that component 1 accounted for 75% of the total conformational change, which was associated with a hinge-like motion. Component 2 accounted for 17% of the total conformational change, which was associated with the overall conformational difference between the PBPs of iDrugSnFRs and the other published betaine and choline binding proteins. The liganded and apo biosensor structures partitioned into two groups (Fig. 1). Component 1 accounted for the Venus-flytrap conformational change and was represented best by the structural alignment of varenicline bound to iNicSnFR3adt and the apo iNicSnFR3adt (Fig. 1- figure supplement 1). As in other betaine or choline binding protein structures in the Protein Data Bank, the liganded conformations differed little from each other, and the conformational change between liganded and apo states related to the Venus-flytrap mechanism. Component 2 accounted for a small fraction of the conformational differences between the native structures and optimized biosensor. This small difference was also observed when the choline-bound iAChSnFRdtbp and *Bacillus subtilis* structures were aligned (RMSD of 1 Å) (Fan, 2020).

### Millisecond timescale microperfusion with iCytSnFR_PM

**Cell culture**. The iCytSnFR_PM sensor was expressed in HEK293T cells because the high level of protein expression provided by this cell line improved the fluorescent signal/noise ratio. The cells were plated at a density of 1.25 x 10^5^ cells per dish on gelatin-coated, glass coverslips attached to the bottom of a 35 mm culture dish and incubated at 37 °C. Twenty-four h after plating, the cells were transfected with iCytSnFR_PM DNA (200 ng per dish) using Lipofectamine 3000 (ThermoFisher Scientific, Waltham, MA). The transfection medium was removed after a 24 h incubation and replaced with fresh growth medium. We incubated the transfected cells for another 24-72 h to obtain sufficient iCytSnFR_PM membrane protein density for fluorescent imaging.

**Fluorescent microscopy.** A blue LED (488 nm wavelength) provided epifluorescent illumination. The bandwidth and peak of the excitation and emission spectra were 460-500 nm and 480 nm (excitation), and 515-550 nm and 535 nm (emission), respectively (cat. #41001 FITC filter cube). Immediately prior to imaging, we removed the culture dishes from the incubator and rinsed them three times with HBSS warmed to 37 °C. After rinsing, we pried the coverslip off the bottom of the dish with a single-edged razor blade and re-sealed it to the bottom of a plastic perfusion chamber (Model RC-25 diamond shape, Warner Instruments) with silicone grease. At this point, the chamber and coverslip were mounted on a motorized microscope stage and perfused continuously with HBSS at 18°C from a gravity-fed, bath perfusion system to keep the cells oxygenated. The fluid level in the chamber was regulated by suction from a syringe needle.

**Drug microperfusion**. The major components of the microperfusion system were (1) a parallel array of three, square glass capillaries (600 µm i.d.) that delivered the ligand to the cells, (2) a stepping motor that translated the capillary array horizontally above the coverslip surface, and (3) a controller that determined the motion and step size of the motor. Both the array and stepping motor were mounted on a motorized micromanipulator (Model MP-285, Sutter Instrument Co. (Novato, CA) that allowed us to position it in the chamber. Fluid flowed continuously from all three capillaries into the chamber during the experiment, as well as vehicle from a separate input connected to the bath perfusion system. The center capillary in the array contained vehicle plus drug, while the two outer capillaries contained vehicle only. Outflow from the end capillaries kept ligand flow from the center capillary confined to a laminar stream. We applied drugs to the cells in 5-30 s step-like pulses. The array was positioned so that outflow from an end capillary bathed the cells in the field of view with vehicle initially. To apply the ligand to the cells, we stepped the entire array horizontally by 700 µm so that vehicle plus drug from the center capillary bathed the cells. Ligand application was terminated by returning the array to its initial position.

**Data acquisition**. A computer equipped with the Clampex v.9 software and a Digidata 1200 series interface (Molecular Devices, San Jose, CA) controlled both the timing of ligand application and initiated image acquisition. A camera (Model ORCA-3G, Hamamatsu (Hamamatsu City, Japan)) attached to a microscope port recorded the images. A second computer equipped with the HCImage V3.0 software and a digital camera interface (Hamamatsu) controlled the camera operation, set the parameters of image acquisition, and stored the recorded image sequences. We recorded image sequences of 170-200 frames at rates of 1-8.9 frames per s to visualize the time course of the iCytSnFR_PM response. Event markers in the image file marked the timing of ligand application. We started acquiring images 5-10 s before applying the ligand to establish a baseline fluorescence and continued for 5-180 s after applying drug to record the decay of the response.

### Millisecond timescale microperfusion with iCytSnFR_PM (Results)

**Biphasic decay components of the cytisine response.** One-way ANOVA showed a significant difference between the relative amplitudes of the decay components after the 5, 8, 10, and 15 µM cytisine pulses (ANOVA, *p* < 0.01, *df* = 3,32). Post-hoc comparisons showed that the relative amplitude of the slow component *A_s_*/(*A_f_* + *A_s_*) (where *A_s_* was the amplitude of the slow component and *A_f_* was the amplitude of the fast) of the 5 μM cytisine decay (24 ± 7%, n = 6 areas (13 cells)) was significantly (Tukey tests, (*p* < 0.05) less than the mean *A_s_*/(*A_f_* + *A_s_*) of the three other concentrations (8, 10, 15 µM) (66 ± 1%, n = 30 areas (92 cells)). The *A_s_*/(*A_f_* + *A_s_*)’s after pulses of 8, 10, and 15 µM cytisine did not differ significantly (*p* > 0.05).

**Growth phase of the cytisine response.** We visualized the time course of the growth phase of the iCytSnFR_PM response to 1-15 µM cytisine, using the Fast-Step microperfusion system (Supp. Figure 2A-B). The growth phases of responses to 30 s applications of 1-4 µM cytisine were biphasic and fitted best by the sum of two terms describing an exponential rise to a maximum (Supp. Figure 2A). The range of the faster rate constants (*kf_on_*’s) for 1-4 µM cytisine was 0.41-1.22 s^-1^ (n = 40 areas (138 cells)) and the range for the slower rate constants was 0.01-0.1 s^-1^ (n = 37 areas (128 cells)). The faster component dominated the rising phase of the 1-4 µM cytisine response. The relative amplitude of the faster component *A_f_*/(*A_f_* + *A_s_*) was 77 ± 1%, n = 37 areas (128 cells)). It was not affected significantly by cytisine concentration in the 1-4 µM range (ANOVA, *df* = 3 model, 33 error, *p* = 0.08). The responses to 5-10 s applications of 5-15 µM cytisine reached steady state within a 5 s application. They were fitted adequately by a single negative exponential rise to a maximum (Supp. Figure 2B). The single rate constant for the rising phase in individual areas ranged from 0.86 to 2.65 s^-1^ (n = 37 areas (133 cells)). We pooled the fast rate constants for the 1-4 µM cytisine responses with the single rate constants for the 5-15 µM responses to obtain an overall [cytisine]-*kf_on_* relation for the 1-15 µM cytisine concentration range (Supp. Fig. 3A). Linear least-squares regression showed that the [cytisine]-*kf_on_* relation over this concentration range was approximately linear (*r* = 0.98, n = 8 concentrations, *p* < 0.05) with a slope and intercept of 1.5 ± 0.1 10^5^ (Ms)^-1^ and 0.43 ± 0.05 s^-1^ (± SE), respectively.

Nevertheless, the [cytisine]-*kf_on_* data from the microperfusion experiments deviated from linearity at cytisine concentrations ≥ 8 µM, suggesting that [cytisine]-*kf_on_* relation for the microperfusion data was more hyperbolic than linear. Using the slower decay rate constant for cytisine (0.146 ± 0.006 s^-1^) as the value for the *kf_on_* at 0 µM cytisine, a hyperbolic relation fitted the cytisine-*kf_on_* relation for 0-10 µM cytisine significantly better than a straight line (Supp. Fig. 3B, F-test, *p* < 0.05).

In contrast to the faster rate constant of the rising phase *kf_on_*, the slower rate constant *ks_on_* decreased significantly as the cytisine concentration increased from 1 to 4 µM (*r* = 0.78, n = 4 concentrations, *p* < 0.05, Supp. Fig. 3D). The slope and intercept of a straight line fit to the [cytisine]-*ks_on_* data using least-squares regression were -1.5 ± 0.06 X 10^4^ M^-1^s^-1^ and 0.09 ± 0.2 s^-1^ (± SE, n = 4 concentrations), respectively.

### Comments on a kinetic model

Our subsecond data with cytisine at iCytSnFR are the most complete, comprising both stopped-flow and HEK293T cell microperfusion. We therefore discuss the cytisine-iCytSnFR kinetics.

Recent literature on kinetics of PBPs emphasizes a modified induced-fit model with the additional possibility that the apo PBP can also undergo spontaneous activation (termed the “closed state” in the SBP literature (de Boer, Gouridis, Muthahari, & Cordes, 2019; Gouridis, et al., 2015). Such a scheme, shown in Supplementary Figure 4, resembles the three-state model we and colleagues developed to account for iSeroSnFR (Unger at al. 2020).

We simulated the scheme using the MATLAB Simbiology toolbox. The following rate constants account for the stopped-flow and millisecond perfusion data within a factor of three: k_bind2_, 0.5 x 10^6^ /M/s; k_unbind2_, 0.4 /s; k_iso(+)_, 0.001 /s; k_iso(-)_, 0.01 /s; the apo fluorescent state has 0.1 times the brightness of the bound fluorescent state.

The three-state model predicts the experimental observation (Supp. Fig. 3) that the rate constant of the slower component of the kinetics decreases as the ligand concentration increases. For the cytisine-iCytSnFR case reported in this paper, we conclude that the apo, fluorescent state is less bright than the bound state (shown by the different colors of the cpGFP moiety).

Interestingly, the three-state model fitted the kinetic data best if we assumed that there is a population of higher-sensitivity iDrugSnFRs in HEK293T cells, with an EC_50_ at least 10 times less than we observed with the stopped-flow and HeLa cell data. A more complex, “square” four-state model, comprising both ligand binding and protein conformational changes, has been applied to equilibrium measurements on cGaMP (Barnett, et al., 2017). Full kinetic predictions of the four-state model are available (Lancet & Pecht, 1976).

The fragmentary kinetic data for acetylcholine at iCytSnFR_PM suggest equilibrium and rate constants in the same broad range as for cytisine. However, the kinetic data for varenicline suggest that ligand unbinding dominates the decay phase, with a rate constant < 0.01 s^-1^. As a caution, recent data show that mechanisms at PBPs (part of the larger class of substrate-binding proteins, SBPs) can change fundamentally with even a single mutation (Nguyen, Lai, Lee, Kaiser, & Rees, 2018). We therefore wish to avoid generalizing past the single iDrugSnFR we consider here.

**Supplementary Figures**

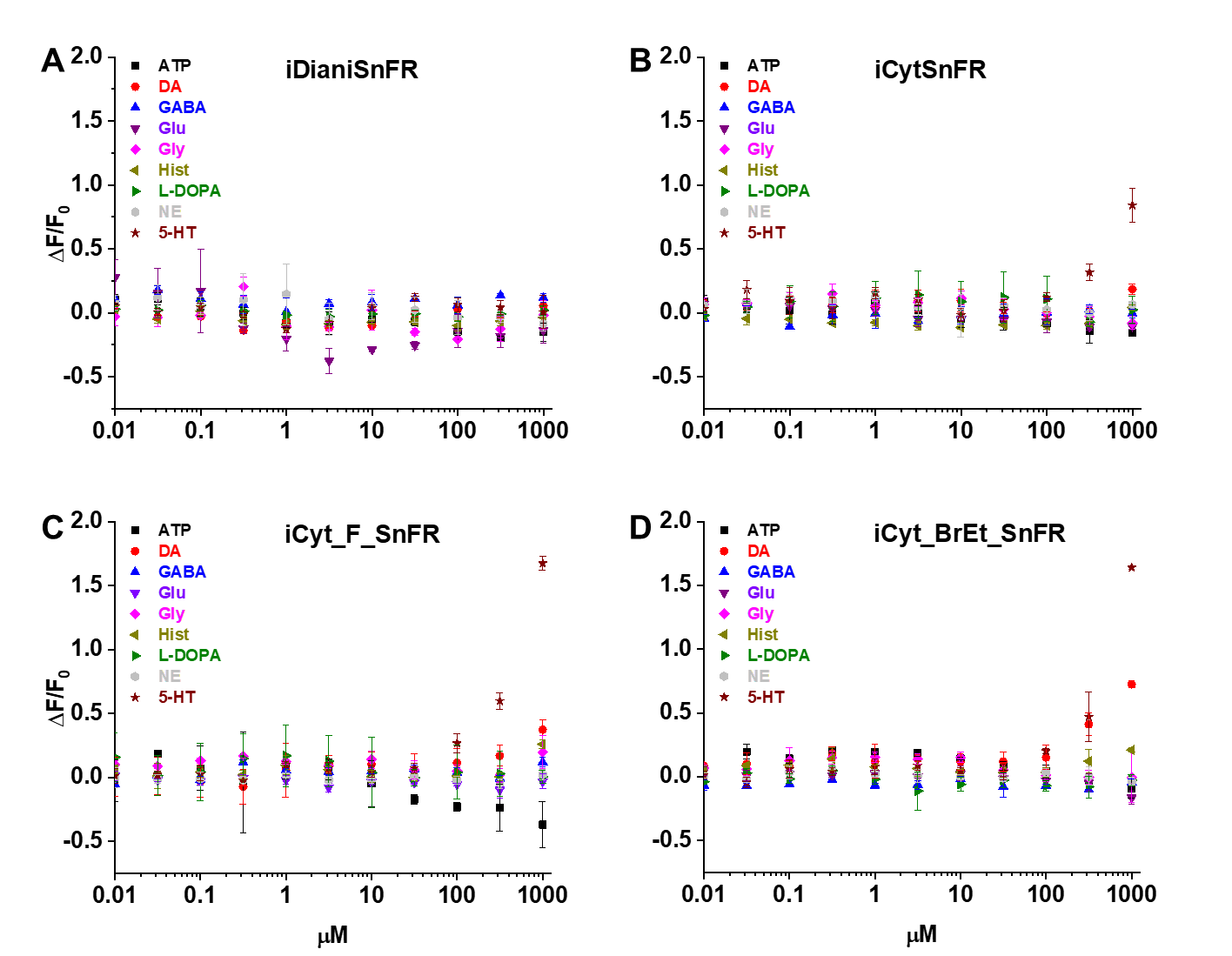

**Supplementary Figure 1**. Dose-response relations of nicotinic agonist iDrugSnFRs against select endogenous molecules. Abbreviations: ATP (adenosine triphosphate), DA (dopamine), GABA (γ-aminobutyric acid), Glu (glutamate), Gly (glycine), Hist (histamine), L-DOPA (levodopa), NE (norepinephrine), and 5-HT (serotonin). **(A)** iDianiSnFR shows no fluorescent response to any of the selected endogenous molecules. **(B)** iCytSnFR, **(C)** iCyt_F_SnFR, and **(D)** iCyt_BrEt_SnFR show no response to any of the selected endogenous molecules except 5-HT and DA at concentrations above 100 µM.

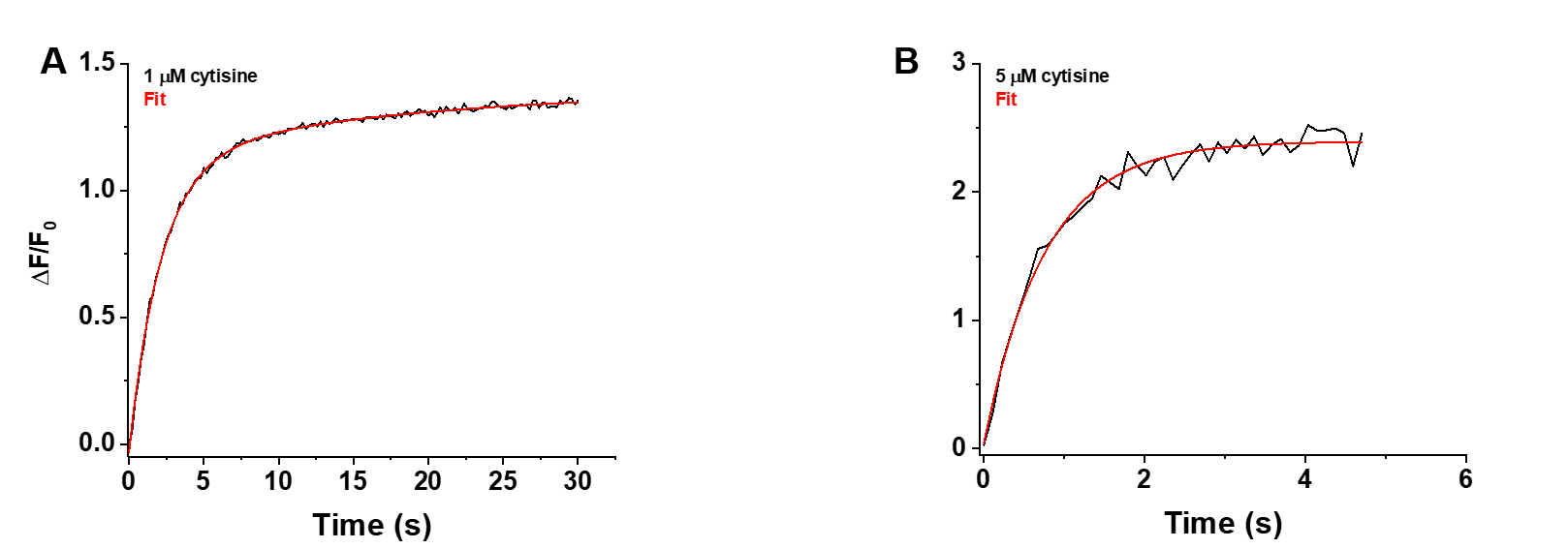

**Supplementary Figure 2.** Rising phase of the iCytSnFR_PM response to cytisine in HEK293T cells. (**A)** Example of the biphasic rising phase of a 1 µM cytisine response in an individual area (black trace, mean of 4 cells). Cytisine was applied for 30 s. The fast and slow time constants of the rising phase (*τf_0n_*, *τs_0n_*) were 2.15 ± 0.05 s and 17 ± 3 s (n = 151 frames, sampling rate of 5 frames/s), respectively. The red line is a fit to the sum of two declining exponentials (*R^2^* of 0.998). It was significantly better than that to a single negative exponential rise to maximum component and a constant term (F-test, *p* < 0.05). The *A_s_/(A_s_+A_f_*) was 20%. **(B)** Example of the rising phase of a response to a 5 s application of 5 µM cytisine in an individual area (mean of 10 cells, 2 replicates). The response appeared to be monophasic with a single time constant (τ_0n_) of 0.76 ± 0.04 s (n = 43 frames, sampling rate of 9.8 frames/s). The red line is a fit to the sum of a negative exponential rise to a maximum, and constant, term (*R^2^* of 0.98).

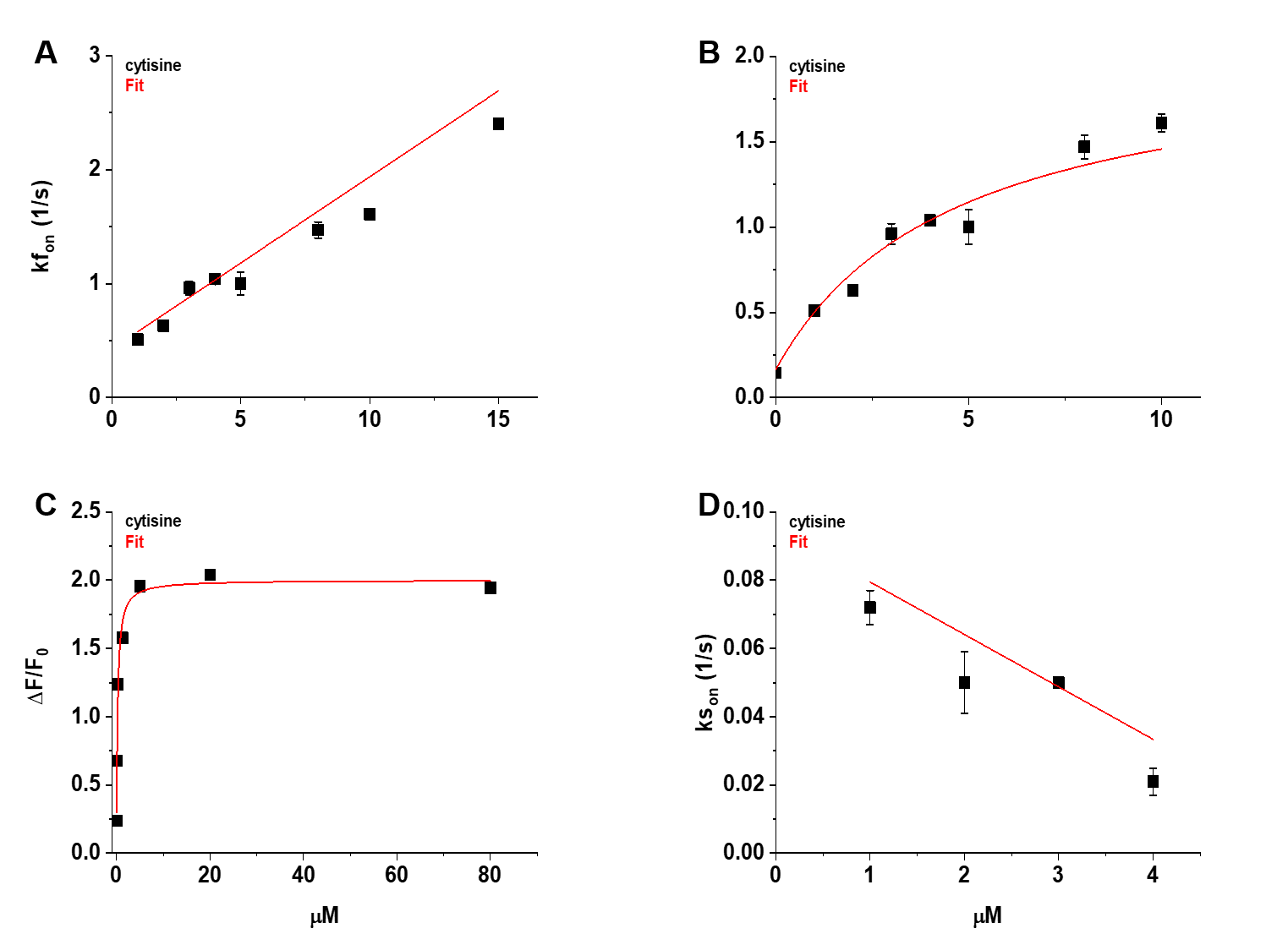

**Supplementary Figure 3.** Concentration dependence of the fast (*kf_on_*) and slow rising rate constants (*ks_on_*) of the cytisine response. **(A)** The [Cytisine]-*kf_on_* relation was approximately linear between 1 and 15 µM cytisine. Symbols (filled squares) are the mean *kf_on_* for the individual cytisine concentrations tested (n = 7-10 areas per concentration, (29-40 cells)). The red line is a regression line fit to the data using linear least-squares regression. See text for the values of the slope, intercept, and correlation coefficient. **(B)** The [Cytisine]-*kf_on_* relation for the *kf_on_* was more hyperbolic than linear between 0 and 10 µM cytisine. Red line is a fit to the sum of a hyperbolic, and constant, term using nonlinear least-squares regression. See text for fitted parameters. We used the mean slow decay rate constant (*ks_off_*) of the cytisine response for the *kf_on_* at 0 µM cytisine (0.146 ± 0.006 s^-1^). **(C)** Concentration-response (CR) relation for the mean steady-state response to cytisine of iCytSnFR_PM sensors expressed in HeLa cells (n = 11 cells, see Fig. 7B). Red line is the fit to the sum of a hyperbolic, and constant, terms using nonlinear least-squares regression. See text for fitted parameters. Symbols (filled squares) are the mean values for the final 10 s of the steady state cytisine response. **(D)** The [Cytisine]-*ks_on_* relation for 1-4 µM cytisine. Red line is a regression line fit to the data using linear least-squares regression. See text for the slope, intercept, and correlation coefficient. Symbols (filled squares) are the mean *ks_on_* for the individual cytisine concentration tested (n = 8-10 areas per concentration, (31-38 cells)). Error bars in panels **(A-D)** are ± SEM. Symbols obscure the bars at some concentrations in panels **(A), (B), and (D)**, and all concentrations in panel **(C)**.

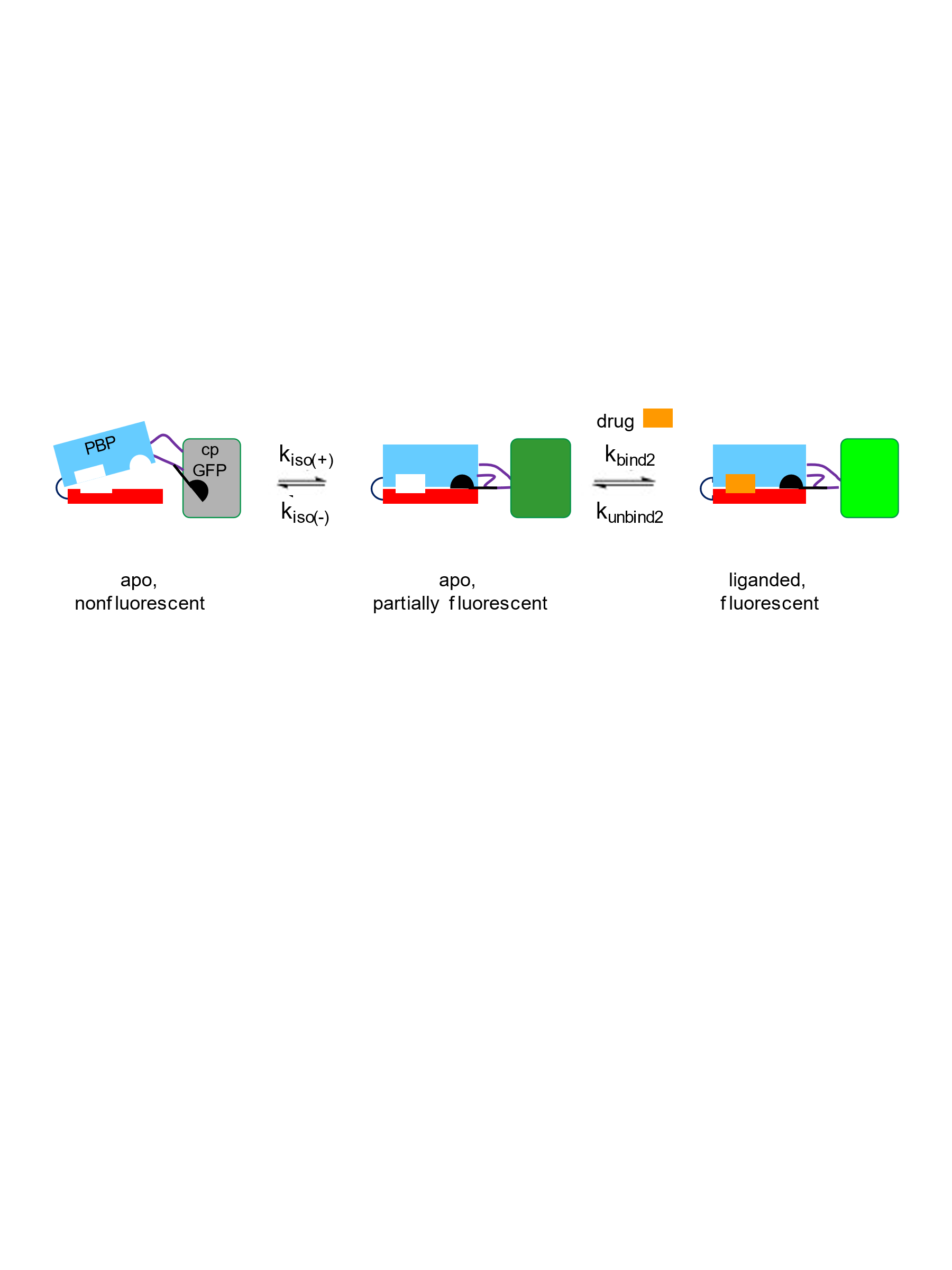

**Supplementary Figure 4**.

A three-state kinetic scheme for iCytSnFR. The diagram contains cartoons of the PBP moiety (blue and red), the linkers (black lines), the Glu78 “candle snuffer” attached to Linker1 (black), and the cpGFP moiety (gray, dark green, or green).

We postulate that the iDrugSnFR exists in an apo nonfluorescent state and an apo fluorescent state; these states interconvert with time constants of tens of seconds k(_iso(+)_, k_iso(-)_). Cytisine binds to the apo fluorescent state (k_bind2_), inducing an additional fluorescent state on a briefer time scale. The initial fluorescence increase represents the binding-induced increase, and the slower increase is governed by partial re-equilibration of the two apo states. Upon removal of cytisine after just a few s of perfusion (Figure 6), the fluorescence decay represents the dissociation of cytisine (k_unbind2_). This scheme resembles the model we and colleagues developed to account for iSeroSnFR (Unger at al. 2020). For the iDrugSnFRs reported in this paper, we conclude that the apo, fluorescent state is less bright than the bound state (shown by the different colors of the cpGFP moiety).

**Supplementary Figure 5.** Traces of fluorescence responses during time-resolved low-concentration dose-response relations for nicotinic agonists in HeLa cells. BC = Buffer control. SEM of data are indicated by semi-transparent shrouds around traces where trace width is exceeded. Cyt (cytisine) in cells expressing iCytSnFR_ER **(A)** or iCytSnFR_PM **(B)**; 10FC (10-fluorocytisine) in cells expressing iCyt_F_SnFR_ER **(A)** or iCyt_F_SnFR_PM **(B)**; 9Br10EtC (9-bromo-10-ethylcytisine) in cells expressing iCyt_BrEt_SnFR_ER **(A)** or iCyt_BrEt_SnFR_PM **(B)**. Relatively long (300 s) washout periods between drug applications allowed a return to baseline fluorescence for the **(A)** ER and **(B)** PM. **(C)** A zoomed in exemplar comparison of the ER and PM for a pulse of 1 µM 10-fluorocytisine shows a distinct lag in the decrease of the fluorescent signal in the ER as compared to the PM.

**Supplementary Tables**

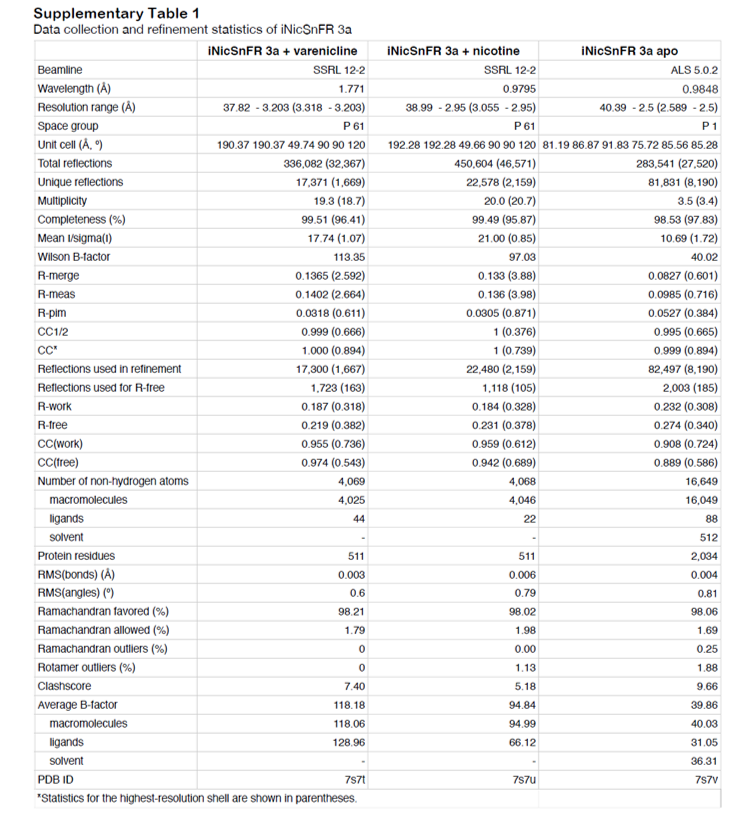
**Supplementary Table 1.**

**Supplementary Table 2**

|  | **iDianiSnFR** | | | **iCytSnFR** | | | **iCyt_F_SnFR** | | | **iCyt_BrEt_SnFR** | | |
| --- | --- | --- | --- | --- | --- | --- | --- | --- | --- | --- | --- | --- |
| **µM** | **t1** | **t2** | **R^2^** | **t1** | **t2** | **R^2^** | **t1** | **t2** | **R^2^** | **t1** | **t2** | **R^2^** |
| **1000** | **0.0067 ± <0.0001** | **0.18 ± <0.01** | **0.986** | **0.027 ± <0.001** | **-** | **0.999** | **0.021 ± <0.001** | **0.32 ± <0.01** | **0.999** | **0.02 ± <0.01** | **-** | **0.992** |
| **500** | **0.0062 ± <0.0001** | **0.22 ± <0.01** | **0.989** | **0.039 ± <0.001** | **-** | **0.999** | **0.027 ± <0.001** | **0.34 ± <0.01** | **0.999** | **0.023 ± <0.001** | **-** | **0.994** |
| **250** | **0.0061 ± <0.0001** | **0.34 ± 0.01** | **0.991** | **0.063 ± <0.001** | **-** | **0.999** | **0.038 ± <0.001** | **0.42 ± <0.01** | **>0.999** | **0.03 ± <0.01** | **-** | **0.997** |
| **125** | **0.0068 ± <0.0001** | **0.39 ± 0.01** | **0.995** | **0.11 ± <0.01** | **-** | **>0.999** | **0.066 ± <0.001** | **0.63 ± <0.01** | **>0.999** | **0.045 ± <0.001** | **-** | **0.998** |
| **31.25** | **0.012 ± <0.0001** | **0.88 ± 0.09** | **0.997** | **0.34 ± <0.01** | **-** | **>0.999** | **0.20 ± <0.01** | **1.7 ± <0.1** | **>0.999** | **0.027 ± <0.001** | **-** | **0.994** |
| **7.8** | **0.025 ± <0.001** | **0.81 ± 0.10** | **0.996** | **0.80 ± <0.01** | **-** | **>0.999** | **0.066 ± 0.014** | **0.99 ± 0.02** | **0.999** | **0.084 ± <0.001** | **-** | **0.998** |
| **1.95** | **0.034 ± <0.001** | **0.44 ± 0.09** | **0.976** | **1.3 ± <0.1** | **-** | **0.994** | **0.14 ± 0.05** | **5.3 ± 4.2** | **0.987** | **0.89 ± <0.01** | **-** | **>0.999** |
| **0.49** | **0.00096 ± 0.00017** | **0.06 ± <0.01** | **0.840** | **0.97 ± 0.06** | **-** | **0.947** | **0.00079 ± 0.00015** | **4.5 ± 2.0** | **0.872** | **1.4 ± <0.1** | **-** | **0.995** |

**Stopped-flow model determined rate constants**

**Supplementary Table 3**

| Video | Description |
| --- | --- |
| Supplementary Video 1 | Video morph of PDB 7S7V to 7S7T |
| Supplementary Video 2 | iDianiSnFR_ER dose-response relations in HeLa cells |
| Supplementary Video 3 | iDianiSnFR_PM dose-response relations in HeLa cells |
| Supplementary Video 4 | iCytSnFR_ER dose-response relations in HeLa cells |
| Supplementary Video 5 | iCytSnFR_PM dose-response relations in HeLa cells |

**Supplementary Videos**
